# Subcellular determinants of orthoflavivirus protease activity

**DOI:** 10.1101/2025.01.31.635871

**Authors:** Lochlain Corliss, Chad M. Petit, Nicholas J. Lennemann

## Abstract

Orthoflaviviruses are small, enveloped, positive-sense RNA viruses that cause over 500 million infections globally each year for which there are no antiviral treatments. The viral protease is an attractive target for therapeutics due to its critical functions throughout infection. Many studies have reported on the structure, function, and importance of orthoflavivirus proteases; However, the molecular determinants for cleavage of intracellular substrates by orthoflavivirus proteases and how these factors affect viral fitness are unknown. In this study, we used our fluorescent, protease-activity reporter system to investigate the subcellular determinants involved in orthoflavivirus protease cleavage. By modifying our reporter platform, we identified endoplasmic reticulum (ER) subdomain localization and membrane proximity of the substrate cut site as two previously uncharacterized molecular determinants for cleavage. We also altered the amino acid composition of the reporter cut site to introduce sequences present at the cytoplasmic junctions within orthoflavivirus polyproteins and found that each protease processed the sequence located at the junction between NS4A and the 2K peptide least efficiently. Live-cell imaging revealed that cleavage of the NS4A|2K sequence is significantly delayed compared to the capsid cleavage sequence. We further determined that introducing a more efficient cleavage sequence into the NS4A|2K junctions of orthoflavivirus infectious clones abolished virus recovery. Overall, this study identifies ER subdomain localization and membrane proximity of the cut site as molecular determinants for cleavage by orthoflavivirus proteases and provides insight into the role that sequence specificity plays in the coordinated processing of the viral polyprotein and establishing productive infections.

**Importance:** Orthoflaviviruses are the most prevalent and dangerous arthropod-borne viruses (arboviruses) leading to over 500 million global infections annually. Orthoflavivirus infection can cause severe pathologies, including hemorrhagic conditions and neurological disease, that lead to hundreds of thousands of deaths each year. The viral protease complex, responsible for processing the viral polyprotein into its functional subunits, is an attractive target for antiviral therapeutic development. Despite extensive research efforts on these viral protein complexes, all protease inhibitor candidates have fallen short of clinical efficacy, highlighting a considerable gap in knowledge of the viral protease’s complex intracellular activity. The significance of our research is in characterizing the subcellular determinants associated with orthoflavivirus protease cleavage efficiency and how these factors can influence viral fitness. These findings contribute to closing this gap in knowledge of the mechanisms of orthoflavivirus proteases which can ultimately lead to the successful development of targeted antivirals.

## Introduction

Orthoflaviviruses belong to a large family of viruses with positive-sense, single-stranded RNA genomes. The viruses in this genus are arthropod-borne and predominantly transmitted by several mosquito species, primarily of the *Aedes* and *Culex* genera (1–4). The arthropod vector of orthoflaviviruses is the main contributor to sporadic epidemic outbreaks in warm geographic locations with high mosquito populations, such as Africa, South and Central America, and Asia (5, 6). However, because of persistent global warming and mosquito migration, previously naïve regions, like the highly populated Gulf Coast and Southwestern coastal region of the United States, are becoming habitable locations for these viral vectors and are at risk of becoming hotspots for outbreaks (7–9). Members of this viral family pose a significant global health concern with dengue virus (DENV) causing an overwhelming ∼ 390 million cases per year (10, 11). Infection with orthoflaviviruses can cause disease leading to severe clinical manifestations in multiple organ systems of the human body. Disease symptoms can include hemorrhagic fever or shock, caused by DENV and yellow fever virus (YFV), encephalitis, caused by West Nile virus (WNV), and birth defects such as microcephaly as a result of Zika virus (ZIKV) infection in pregnant women (12–14). Orthoflavivirus epidemics have devastating effects on human health as well as significant social and economic burdens to the affected regions (15–17). There are few safe and effective vaccines and no approved antiviral treatments for diseases caused by orthoflaviviruses (18–20). Given the worldwide prevalence of orthoflavivirus infections, it is crucial to gain a more in-depth understanding of the molecular processes of these pathogens to aid in the development of effective preventions and treatments.

Upon viral entry and uncoating, the flavivirus genome is translated at the host cell endoplasmic reticulum (ER) into a single polyprotein that is associated with the ER through multiple transmembrane domains (21, 22). The polyprotein is then processed by viral and host proteases into ten functional subunits, consisting of three structural proteins (C, prM, E) and seven non-structural (NS) proteins (NS1, NS2A, NS2B, NS3, NS4A, NS4B, NS5) (23, 24). The viral serine protease, a complex between NS2B and NS3 (NS2B3), facilitates all the cytoplasmic cleavage events of the polyprotein (25–27). NS3 is a soluble, cytoplasmic protein consisting of a protease and helicase domain; However, it remains localized at the ER through its association with NS2B (28). NS2B has multiple transmembrane domains that anchor it to the ER while NS3 associates with the cytoplasmic loop of NS2B. This hydrophilic domain of NS2B is known to serve as a conserved cofactor that is required for the catalytic activity of the protease (29–31). Previous studies have identified a conserved substrate recognition motif based on the sequences processed within the orthoflavivirus polyprotein, and *in vitro* biochemical assays with a truncated recombinant DENV protease have shown this enzyme cleaves a broad range of sequences within the substrate, which contrasts with the WNV protease (26, 32, 33). However, there are limited studies investigating the intracellular activity of various NS2B3 protein complexes and how this contributes to infection.

Given the prominent role of the virally-encoded protease in the virus lifecycle and disease pathogenesis, it has long been an attractive target for viral inhibitors (34–36). To date, all antiviral therapeutics developed against the flavivirus protease that show promising pre-clinical efficacy have been unsuccessful in clinical testing (37). Many inhibitors have been found to only be effective in biochemical assays, suggesting a gap in knowledge of the complex molecular activities of this viral protein in infected cells (38). Thus, a deeper understanding of the mechanism of action behind NS2B3-mediated cleavage during infection is vital for the design and development of effective antivirals targeting this essential viral protease complex.

## Materials and Methods

### Cells and Viruses

Human embryonic kidney-293T cells (ATCC, CRL-3216) and U2OS osteosarcoma cells (ATCC, HTB-96) were cultured in Dulbecco’s modified Eagle’s medium (DMEM) supplemented with 10% fetal bovine serum (FBS) and 100 U/mL penicillin/streptomycin (P/S). Vero E6 cells (a gift from Kevin Harrod, University of Alabama at Birmingham) were cultured in modified Eagle’s medium (MEM) supplemented with 10% FBS and 100 U/mL P/S. C6/36 Aedes albopictus cells (ATCC, CRL-1660) were cultured in DMEM supplemented with 10% FBS and 100 U/mL P/S. All mammalian cells were maintained in a humidified environment at 37C. C6/36 cells were maintained in a humidified environment at 28C.

DENV serotype 2 strain 16681 and ZIKV-MR766 were rescued from HEK 293T cells transfected with pcDNA6.2 DENV2 16681 and pcDNA6.2 ZIKV MR766 plasmids, respectively, as described previously (39, 40). Supernatants were then passaged over C6/36 cells to propagate the virus. Supernatants were collected and clarified to remove cell debris by centrifugation at 2300 xg for 15 minutes at 4C. Clarified supernatants were aliquoted and stored at -80C.

### Plasmid construction

The WT FlavER construct was generated as previously described (41).

#### Membrane proximity reporters

Cloning of the FlavER reporters with extended linker regions was achieved through site-directed mutagenesis. Site-specific forward and reverse primers were used with T7_F (5’-TAATACGACTCACTATAGGG-3’) and FlavER-StuI_R (5’-CGGGAGCTTTTTGCAAAAGCC-3’) primers to generate overlapping PCR fragments that were ligated into the FlavER vector linearized with NheI and StuI via HiFi assembly (NEB). Site-specific primer pair sequences are shown in Table 1. Ligations were transformed into DH5α *E. coli* (Zymo Research).

**Table 1.**
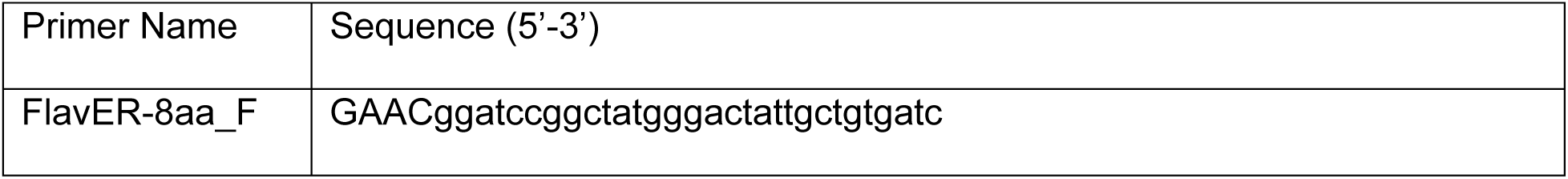
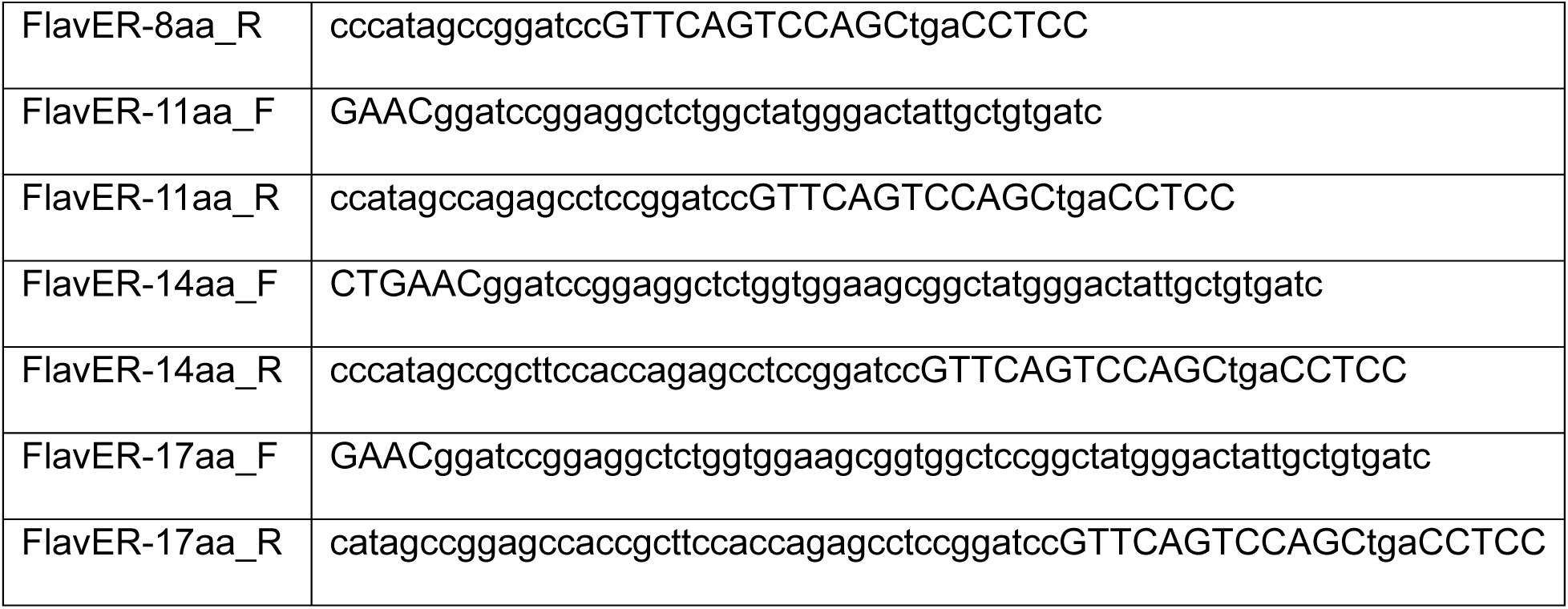

#### ER subdomain reporters

Tubule-Rep was constructed through amplification of Rtn4a (a gift from Carolyn Coyne, Duke University), using Rtn4a-Rep_F and Rtn4a-Rep_R. The protease recognition motif, NLS, and GFP were amplified from FlavER using Rtn4a-RRS_F and GFP-NLS_R. The two fragments were HiFi assembled into the FlavER backbone digested with NheI and ApaI. Sheet-Rep was constructed through amplification of SigmaR1 from the cDNA of HEK293T cells using SigmaR1-Rep_F and SigmaR1-Rep_R. This Product was HiFi assembled with FlavER digested with BamHI and ApaI. Ligations were transformed into DH5α *E. coli*.

**Table 2.**
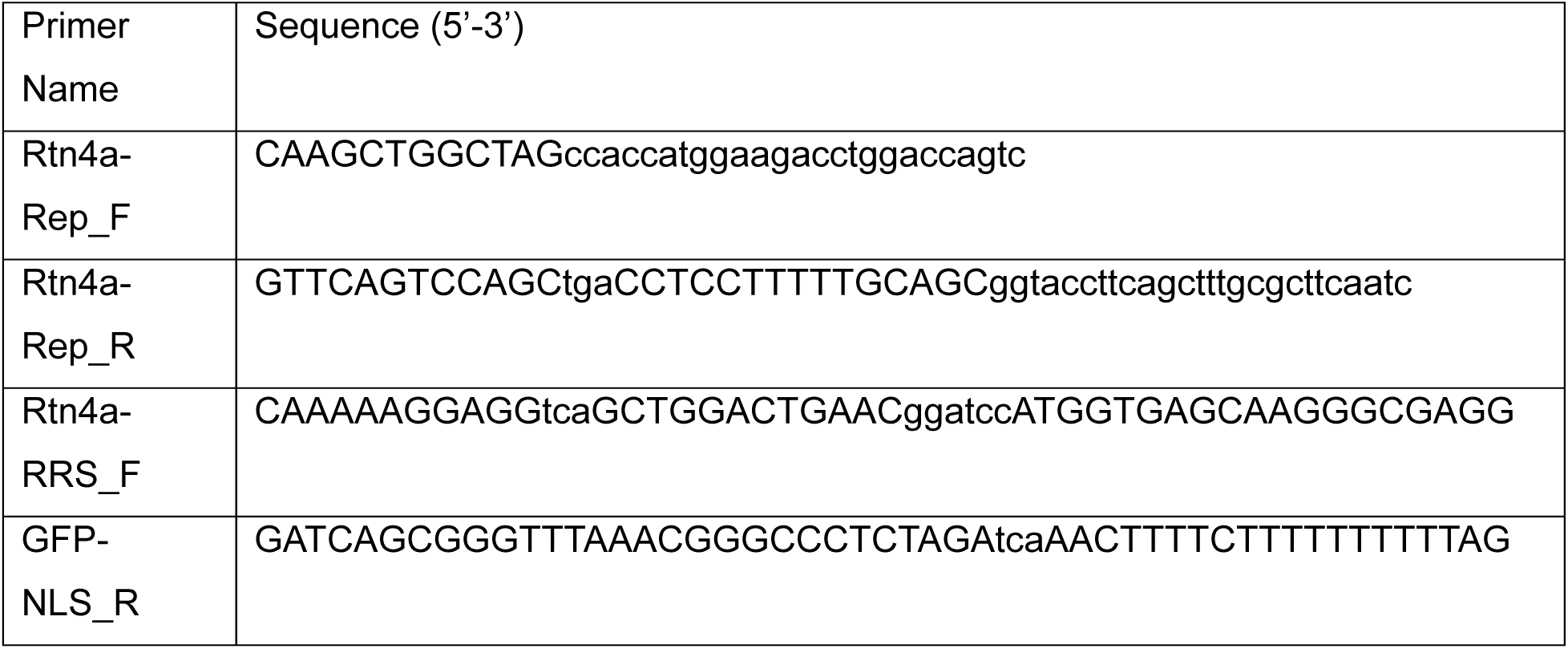
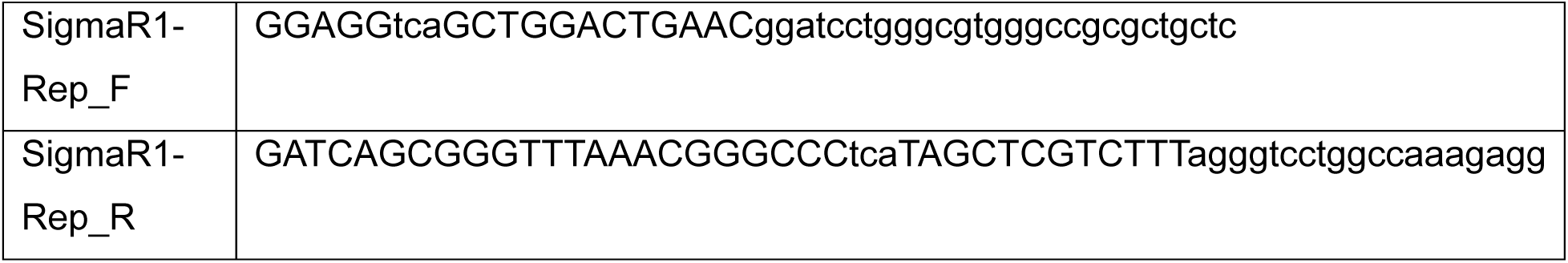

#### Polyprotein junction sequence reporters

Cloning of the FlavER reporters with varying cut site sequences was accomplished through standard restriction enzyme cloning. Sequence-specific oligos were annealed and ligated into the KpnI and BamHI linearized FlavER vector using T4 DNA ligase (NEB) for 30 minutes at room temperature. Annealed oligo sequence pairs are shown in Table 3. Ligations were transformed into DH5α *E. coli*.

**Table 3.**
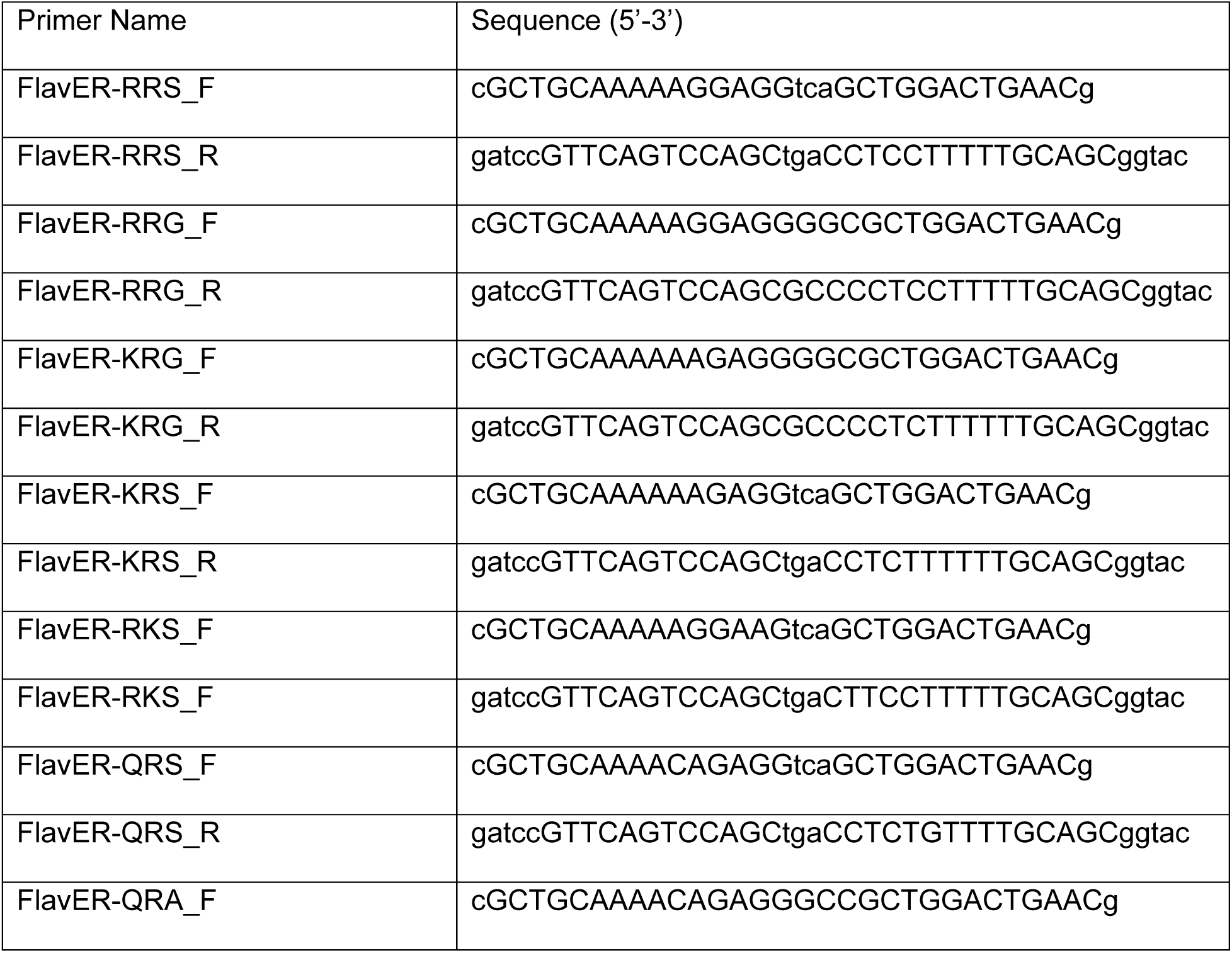
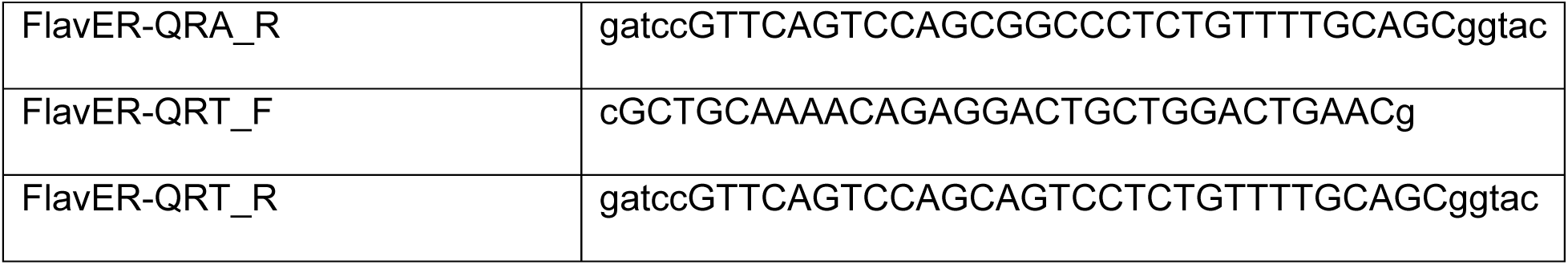

#### Live-cell imaging reporters

Generation of the modified reporters used for live-cell imaging was performed using standard restriction cloning. The mCherry was removed from the FlavER-RR|S and QR|T vectors through digestion with EcoRI and ApaI and replaced with a NanoLuciferase that was synthesized as a gBlock with homology to the reporter vector and ligated using T4 DNA ligase for 30 minutes at room temperature. The GFP from GFP-QR|T-NLuc was cut out of the vector using NheI and XhoI and replaced with the mCherry digested from pcDNA3.1(+)-mCherry using the same restriction enzymes. The two products were ligated using T4 DNA ligase for 30 minutes at room temperature. Ligations were transformed into Stable Competent *E. coli* (NEB).

#### Viral proteases

The construction of expression plasmids encoding DENV NS2B-3-V5, DENV NS2B-3-S135A-V5, ZIKV NS2B-3-V5, WNV NS2B-3-V5, and YFV NS2B-3-V5 have been described previously (41, 42).

#### Site-directed mutagenesis of infectious clones

Mutations to the NS4A|2K cleavage junctions of pcDNA6.2 DENV2 16681 and pcDNA6.2 ZIKV MR766 plasmids were introduced through site-directed mutagenesis. Site-specific forward and reverse primers were used with DENV2-NS3-XhoI_F (5’-GTATAGCAGCTAGAGGATACATCTC-3’) and DENV2-NS5-StuI_R (5’-CTTCGTGTCCTGGTCCTCCTTTTG-3’) primers for pcDNA6.2 DENV2 16681 and ZIKV-NS3-SbfI_F (5’-AGCAGTTGCTCTGGACTACCC-3’) and ZIKV-NS5-MluI_R (5’-GATAAACTCTTCTTTGGTGCAG-3’) primers for pcDNA6.2 ZIKV MR766 to generate overlapping PCR fragments that were ligated into the pcDNA6.2 DENV2 16681 or pcDNA6.2 ZIKV MR766 backbone linearized with XhoI and StuI or SbfI and MluI, respectively, via HiFi assembly (NEB). Site-specific primer pair sequences are shown in Table 4. Ligations were transformed into Stbl2 Competent *E. coli* (Thermofisher).

**Table 4.**
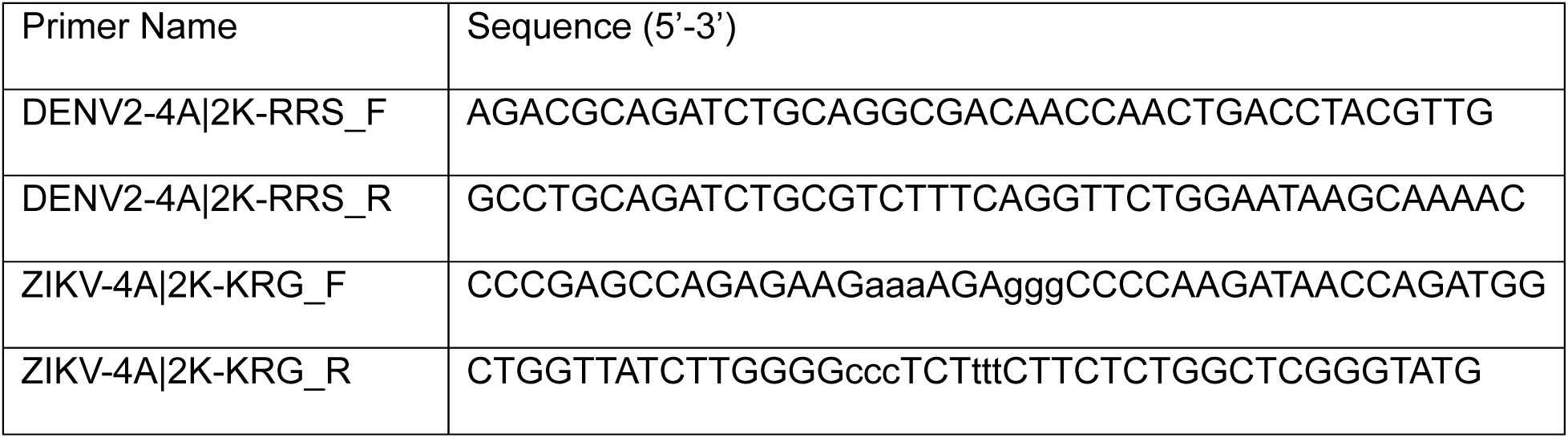

### Molecular modeling

The three-dimensional structure of the NS2B-NS3-NS4A polyprotein (NS^2B-4A^) was generated using Alphafold3 (43). Specifically, we modeled residues 1341-2220 to ensure that NS3 was adequately tethered to the membrane by NS2B and NS4A of the polyprotein. This structure was then modeled into a POPC membrane using VMD (Visual Molecular Dynamics) program (44). Briefly, NS^2B-4A^ was solvated using Helmut Gubmuller’s SOLVATE program (https://www.mpinat.mpg.de/grubmueller/solvate). The POPC membrane and NS^2B-4A^ were then aligned using their center of mass and NS^2B-4A^ was then incorporated into the membrane. The entire system was then solvated and neutralized by adding the appropriate number of NA^+^ ions using VMD. We then used NAMD to perform molecular dynamics (MD) simulations to ensure the model generated was appropriately equilibrated and minimized (45). We used CHARMM36 force-field parameters for all steps of this process (46). Briefly, we first performed minimization and equilibration on the system with all atoms constrained except the lipid tails to appropriately relax the lipid bilayer. We then repeated minimization and equilibration with only the protein constrained to appropriately relax the environment. Finally, the entire system was equilibrated using a short simulation (0.5 ns) at constant temperature (300K) and pressure (1 atm).

### Lentivirus production and transduction

The production of lentiviral vectors expressing the indicated reporter transgenes was performed using the same method as described previously (41). Briefly, (1.5×10^6^) HEK-293T cells were transfected with 1 ug pLJMI-reporter, 0.75 ug psPAX2 (Addgene plasmid #12260), and 0.25 ug pCAGGS-VSV G-Kan using PEI. 48 hours post transfection, lentivirus was harvested by passing cell supernatants through 0.2-micron filters (Thermo Scientific, 725-2520). U2OS cells (5×10^5^) were transduced with 500 uL of each lentiviral vector stock for 48 hours prior to re-seeding into 8-well chamber or live-imaging slides.

### Immunofluorescence staining/microscopy

U2OS cells (40,000 cells/well) stably expressing reporter constructs were infected with DENV or reverse transfected with the indicated V5-tagged DENV protease in an 8-well chamber slide (Celltreat). After 24 hours, cells were fixed in 4% PFA in PBS, permeabilized with 0.1% Triton X-100 (Fisher) diluted in PBS, and washed with PBS followed by incubation with mouse α-V5 epitope tag monoclonal antibody (Invitrogen, 46-0705) or recombinant α-dsRNA monoclonal antibody (Kerafast) for 1 hour at RT. The cells were then washed with PBS 3X for 5 minutes each and probed for 30 minutes with goat α-mouse conjugated to Alexa Fluor-647 (Invitrogen, A21236) at RT. Following secondary antibody incubation, cells underwent 3 PBS washes for 5 minutes each prior to incubation in PBS containing 300 nM DAPI for 5 minutes at RT. The slide was mounted using ProLong Diamond Antifade Mountant (Invitrogen) and a 24×50mm Premium Superslip (FisherScientific). Cells were imaged on the 60X objective of an Olympus IX83 inverted fluorescent microscope.

### Intracellular cleavage assay

U2OS cells (200,000 cells/well) were transfected with the indicated reporter and protease plasmids using polyethyleneimine (PEI) at a 1:1 ratio of DNA (ug) to 1 mg/mL PEI stock (uL). 24 hours post transfection, cells were lysed in 1× RIPA + protease inhibitor cocktail (Sigma). Lysates were clarified by centrifugation at 12,000 xg for 10 minutes prior to separation by SDS-PAGE using a 4-20% Tris-glycine polyacrylamide pre-cast gel (BioRad) and transfer to nitrocellulose membranes. After 30 minutes of blocking in PBS + 10% non-fat milk, membranes were probed using the indicated primary antibodies: mouse α-V5 (Invitrogen), rabbit α-GFP (Proteintech), mouse α-GFP (Proteintech), mouse α-actin (Proteintech), and/or rabbit α-DENV NS3 (GeneTex) followed by incubation with corresponding near-infrared dye conjugated secondary antibodies (LiCor) diluted in PBST + 5% nonfat milk and imaged on an Odyssey CLx imaging system (LiCor).

### Long term fluorescent live-cell imaging

3×10^4^ U2OS cells co-expressing two reporter constructs (GFP-QRT and mCherry-QRT or GFP-RRS and mCherry-QRT) were seeded in an 8-well live-imaging slide (Ibidi). The following day, cells were infected with DENV at an MOI of 10. Infection was synchronized at 4C for 1 hour followed by shifting cells to 37C for 3 hours. 4 hours post infection, the imaging slide was transferred to a CO2-controlled incubated chamber positioned over a motorized inverted microscope. Images were taken every 20 minutes for a 20-hour time period, ending at 24 HPI. Image series were exported and compiled using ImageJ software (NIH, Bethesda, MA, USA) and multi-panel images were cropped and rendered in Photoshop CC 2021 (Adobe, San Jose, CA, USA).

### Focus forming assay

HEK-293T cells were transfected with the WT or mutated DENV or ZIKV infectious clone plasmids. 72 hours post transfection, supernatants were collected and serially diluted over Vero E6 cells for 48 hours. Cells were fixed in 4% paraformaldehyde (Electron Microscopy Sciences) diluted in phosphate buffered saline (PBS) for 10 minutes at room temperature (RT). Cells were then permeabilized with PBS + 0.1% Triton 100-X for 10 minutes at RT followed by a 5-minute wash with PBS. Monolayers were incubated with α-orthoflavivirus E-protein monoclonal antibody (clone 4G2, ATCC VR-1852) overnight at 4C. Cells were then washed with PBS followed by incubation with α-mouse antibody conjugated to Alexa Fluor 488 (Invitrogen A11029) for 30 minutes at RT. Foci were counted using an Olympus IX83 inverted fluorescent microscope.

### Image and data analysis

Models were generated using Photoshop, Adobe Illustrator, and Biorender. Immunoblots were uploaded and quantified in ImageStudio. Quantitative values were determined by dividing the cleavage band by the total reporter signal followed by normalization to protease expression. Data was plotted in Prism 9 Software (Graphpad).

Intensity plot profile analysis was performed using the Plot Profile tool in ImageJ. Data was plotted in Prism 9 Software (Graphpad).

ImageJ was used to quantify fluorescent signal intensity in the nucleus of cells from live-cell time-lapse images. Data was plotted in Prism 9 Software (Graphpad).

### Statistics

Non-linear regression analysis, unpaired t-tests, and one-way ANOVA were performed in Prism 9 Software (Graphpad).

## Results

### Intracellular reporter system for assessing substrate cleavage efficiency by orthoflavivirus proteases

To study flavivirus protease specificity for intracellular substrates, we utilized our recently developed protease-dependent reporter system (41). This modified reporter construct is designed with a green fluorescent protein (GFP) fused to a nuclear localization signal (NLS) followed by an interchangeable 10 amino acid protease recognition motif. The recognition motif is then fused to a transferrin receptor transmembrane domain (TM) that anchors a red fluorescent protein (mCherry) into the lumen of the ER (Fig. 1A). In cells co-expressing our reporter with a catalytically inactive DENV protease (Dpro-S135A), both fluorescent signals stay co-localized at the ER; however, in cells co-expressing our reporter with an active DENV protease (Dpro), we observe nuclear translocation of the GFP signal indicating cleavage of the recognition motif while the mCherry signal remains localized to the ER (Fig. 1C). Similarly, we observe the same phenotype of nuclear translocation of GFP and ER-retention of mCherry in reporter-expressing cells that are infected with DENV (Fig. 1D). Probing the lysates of cells transfected under the same conditions as Figure 1C revealed a ∼30 kDa cleavage band (GFP-NLS) from cells expressing Dpro, while this cleavage band was absent in cells expressing Dpro-S135A (Fig. 1B). The same cleavage fragment is observed from the lysates of reporter-expressing cells that are infected with multiple flaviviruses (41). Together, these results indicate that reporter cleavage can be observed visually using immunofluorescent staining or immunoblotting methods.

**Figure 1.**
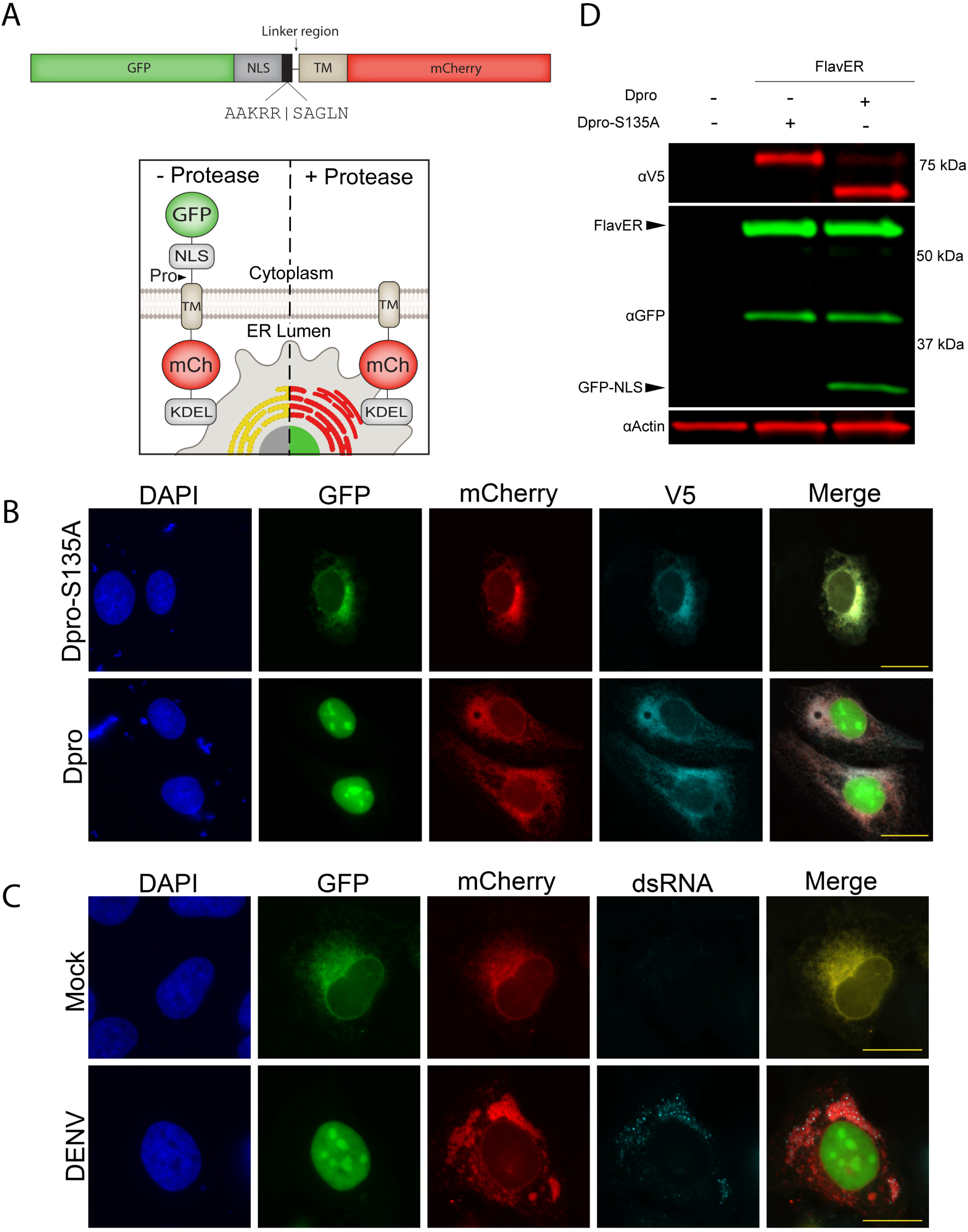
Design and validation of flavivirus protease activity reporter (FlavER) A.) Linear and cartoon model of FlavER. (A, top panel) Linear model of FlavER. The WT FlavER reporter consists of a GFP fused to an NLS followed by a consensus viral protease cleavage motif that is fused to a TM domain followed by an mCherry fluorescent protein. Black box represents the sequence cleaved by the viral protease and black line represents the linker region between the cleavage motif and the TM. (A, bottom panel) Cartoon model of FlavER function. In cells lacking viral protease expression (left), the GFP and mCherry signals co-localize at the ER. In cells expressing a flavivirus protease (right), the NS2B/NS3pro recognition sequence is cleaved, allowing for nuclear translocation of the reporter signal. B.) Immunofluorescence (IF) microscopy of FlavER-expressing U2OS cells transfected with WT DENV protease (Dpro) or catalytically inactive DENV protease (Dpro-S135A). GFP signal is shown in green, mCherry is shown in red, V5 staining is shown in cyan, and nuclei are stained with DAPI shown in blue. Scale bars represent 20 um. C.) IF microscopy of FlavER-expressing U2OS cells infected with DENV at an MOI of 10. GFP signal is shown in green, mCherry is shown in red, dsRNA staining is shown in cyan, and nuclei are stained with DAPI shown in blue. Scale bars represent 20 um. D.) Immunoblot for V5, GFP, and actin of U2OS cells co-transfected with FlavER and the indicated DENV protease. Labeled arrows denote the full-length (FlavER) and cleaved (GFP-NLS) reporter construct. Dpro-S135A represents the catalytically inactive DENV protease. **\***, denotes an intermediate product caused by non-specific proteolysis of mCherry as previously reported (41, 97).

### ER membrane proximity as a molecular determinant of DENV protease cleavage

We sought to use our protease activity reporter system to assess the potential molecular determinants involved in cleavage specificity. During our validation of the FlavER reporter, we found that the construct must have a TM domain that anchors it to the ER membrane in order to be cleaved by Dpro, suggesting that membrane association of intracellular substrates is an important factor for protease specificity (41). These results led us to hypothesize that the proximity of the substrate’s cut site to the ER membrane may serve as a cleavage determinant. This hypothesis is supported by molecular modeling, which shows the catalytic serine of the ER-tethered protease domain is 16.8 Å from the membrane, suggesting that the protease is highly restricted to substrates present at the ER membrane (Fig. 2A). Consistently, the predicted distances of the majority of cleavage junctions within the DENV polyprotein are positioned less than 10 amino acids away from the closest TM domain, suggesting the potential for an optimal membrane distance of a substrate’s cleavage site (Fig. 2B). In order to determine if cut site membrane proximity is involved in cleavage efficiency, we modified our WT FlavER to increase the number of amino acids in the linker region between the protease recognition site and the TM domain from 8 to 17 residues in increments of 3 (Fig. 2C). These constructs were then used in intracellular cleavage assays where we co-expressed Dpro with a reporter encoding a membrane proximity of 8, 11, 14, or 17 residues and probed the cell lysates for GFP to reveal the ratio of full-length to cleaved reporter and the V5 epitope tag to determine the expression of the viral protease. These experiments resulted in a consistent and proportionate decrease in reporter cleavage efficiency as the cut site’s membrane distance increased from 8 to 17 residues (Fig. 2D, E). These results were further recapitulated in the context of infection when cells expressing the membrane proximity reporters were infected with DENV for 24 hours (Fig. 2F). These data suggest that membrane proximity of a substrate’s protease recognition site is involved in the efficiency of substrate cleavage and that there is an inverse correlation between increased distance and cleavage efficiency.

**Figure 2.**
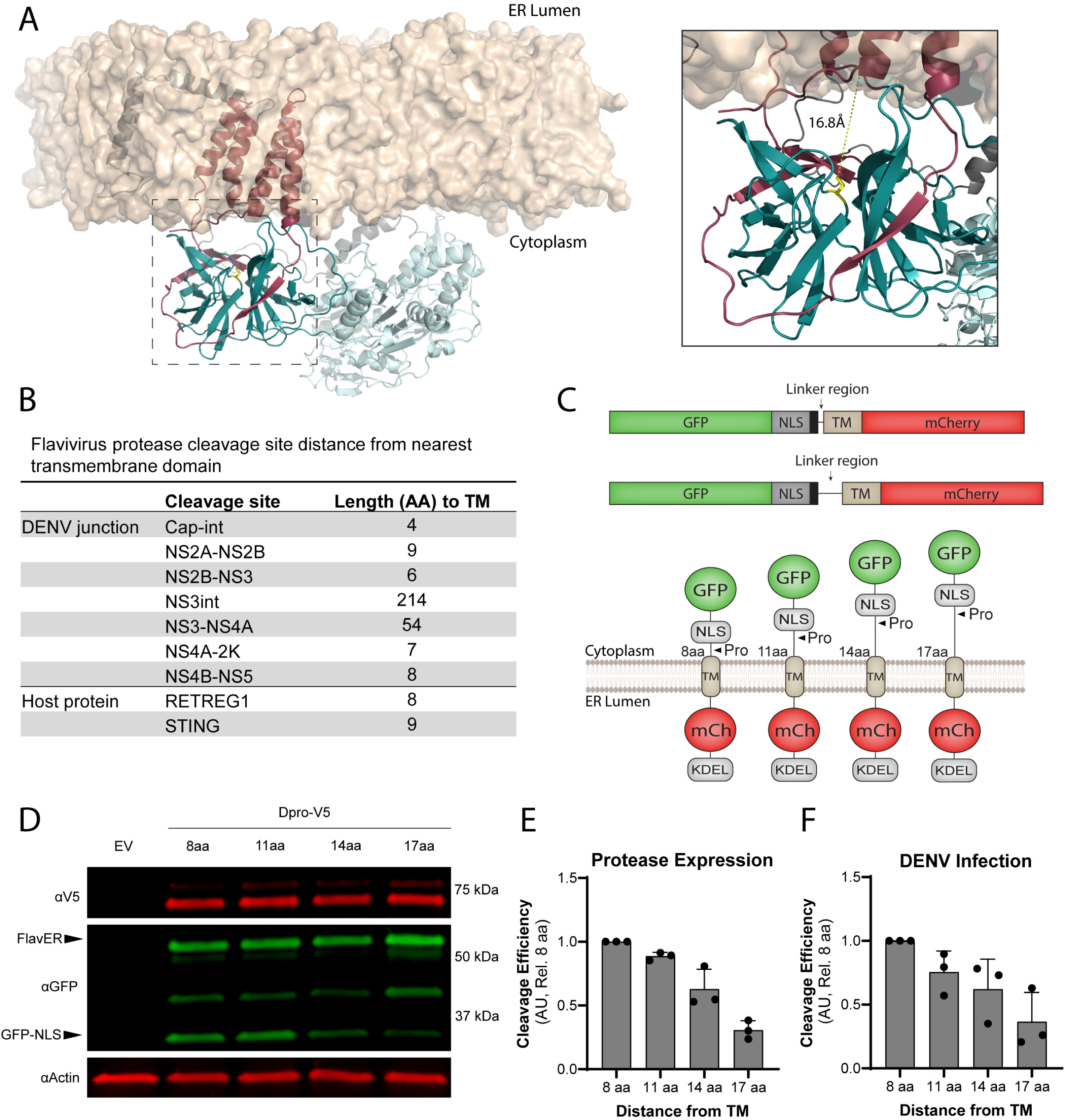
Membrane proximity of substrate cut site as a molecular determinant for cleavage. A.) *Left*, molecular model of the NS2B3 complex in relation to a lipid bilayer. *Right*, zoom-in of boxed region. NS2B is shown in dark pink, NS3 protease domain is shown in dark cyan, catalytic serine shown in yellow, NS3 helicase domain is shown in light cyan, NS4A shown in gray, and lipid bilayer is shown in tan. Distance from the catalytic serine to the membrane represented as yellow line (16.8 Å). B.) Table showing the predicted distances of the protease recognition site from the nearest TM domain within the cytoplasmic cleavage junctions of flavivirus polyproteins. C.) Linear and cartoon models representing the reporter constructs with lengthened linker regions between the protease recognition site and the TM domain. D.) Immunoblot for V5, GFP, and actin of U2OS cells co-transfected with the indicated reporter construct and Dpro. Labeled arrows denote the full-length (FlavER) and cleaved (GFP-NLS) reporter construct. E-F.) Quantification of cleavage assays from reporter-expressing cells transfected with Dpro (D) or infected with DENV at an MOI of 3 (E). Data are presented as efficiency of cleavage determined by intensity ratios in arbitrary units (AU) of the cleaved to total reporter signal relative to viral protease expression. Quantification represented as cleavage efficiency relative to the FlavER-8aa construct. Each bar represents the average cleavage efficiency of three experiments per condition and black circles indicate the cleavage efficiency calculated for each replicate.

### ER subdomain localization as a molecular determinant DENV protease cleavage

Although the ER is a single, interconnected network composed of a contiguous membrane, this dynamic organelle is known to have functionally and structurally distinct subdomains consisting of curved tubules and flat sheets (47–51). Because the established relationship between the ER and flavivirus replication cycle has been well documented, we next wanted to investigate the effects of ER subdomain localization of the substrate on cleavage efficiency by the DENV protease (52–59). To examine if the DENV protease has a specificity for cleavage of substrates localized to ER tubules versus sheets, we modified our reporter system to fuse a fluorescent protein, NLS, and protease recognition motif to host proteins known to localize to these different subdomains of the ER. For the ER tubule construct, our reporter motif (GFP-NLS-cut site) was fused to the C-terminus of reticulon-4a (Rtn4a), while the sigma 1 receptor (SigmaR1) was inserted downstream of the reporter motif for the ER sheet construct to ensure proper topology and membrane proximity of the cleavage site (Fig. 3A, B). We co-expressed the constructs in cells together and observed by fluorescence microscopy and plot profile analysis that although both reporters reside in the same organelle, their respective fluorescent signals are localized to the distinct subdomain to which they are targeted (Fig. 3C, D). These reporter constructs were used in the same intracellular cleavage assays as described previously with both Dpro overexpression and DENV infection. We observed that the DENV protease had higher cleavage efficiency of the ER sheet reporter compared to our WT and ER tubule reporter constructs in both conditions, despite having lower protease expression in Sheet-Rep samples (Fig. 4A-E). Interestingly, we found that even though the DENV protease was able to cleave the ER tubule reporter during co-expression, the cleavage fragment was absent in the assays performed with cells infected with DENV, suggesting that the DENV protease is unable to cleave substrates specifically localized to ER tubules during infection (Fig. 4C, E). Together, these results indicate that the DENV protease has a strong preference for substrates localized to ER sheets.

**Figure 3.**
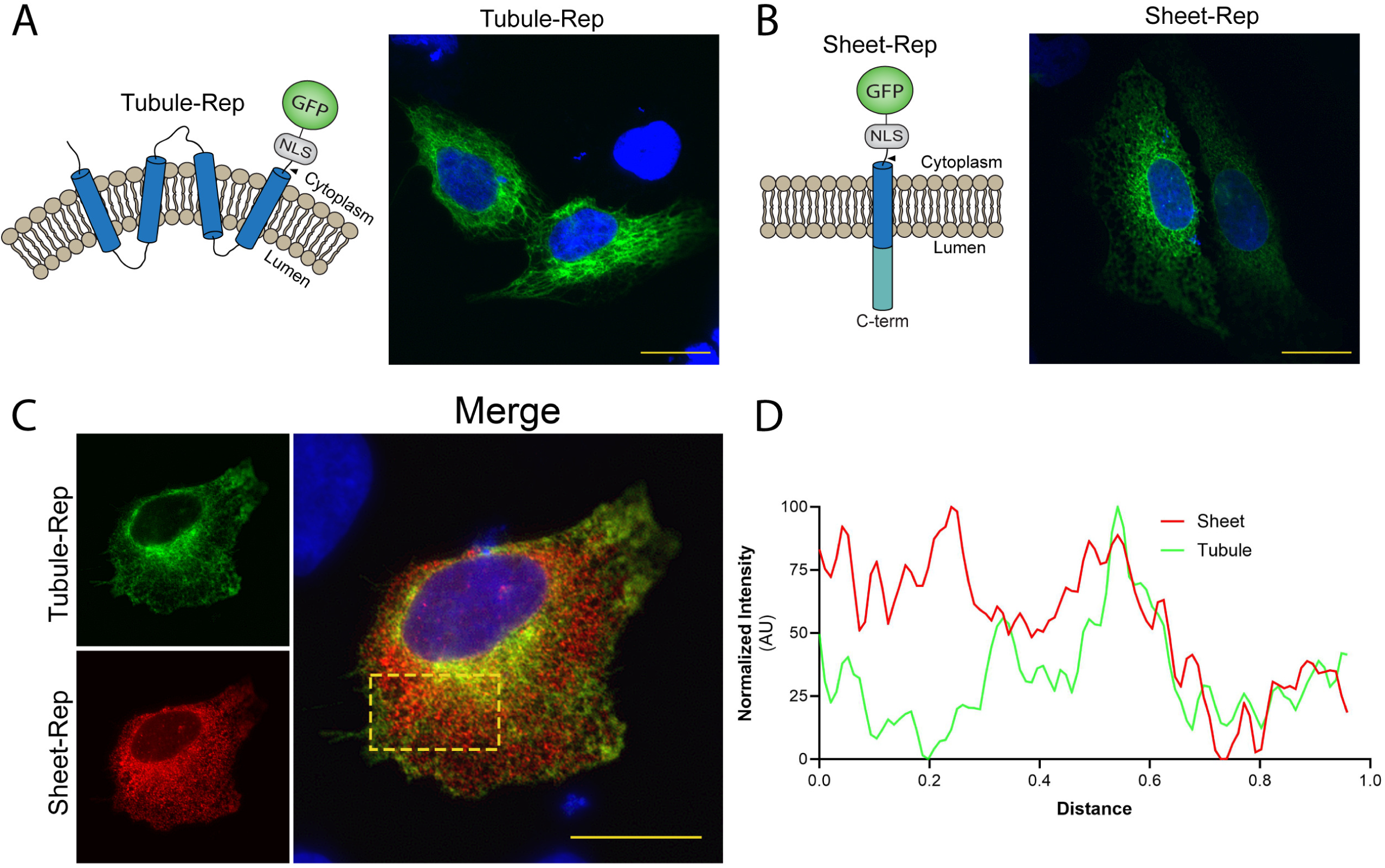
Development of reporters localized to specific ER subdomains. A-B.) Cartoon model and representative IF microscopy images of U2OS cells expressing GFP-Tubule-Rep (A) and GFP-Sheet-Rep (B). GFP signal is shown in green, and nuclei are stained with DAPI and shown in blue. Scale bars represent 20 um. C.) IF microscopy of U2OS cells co-transfected with GFP-Tubule-Rep and mCh-Sheet-Rep. GFP-Tubule-Rep shown in green, mCh-Sheet-Rep shown in red, and nuclei are stained with DAPI and shown in blue. Scale bars represent 20 um. D.) Plot profile of normalized signal intensities from representative GFP-Tubule-Rep (green) and mCherry-Sheet-Rep (red) expressing cells.

**Figure 4.**
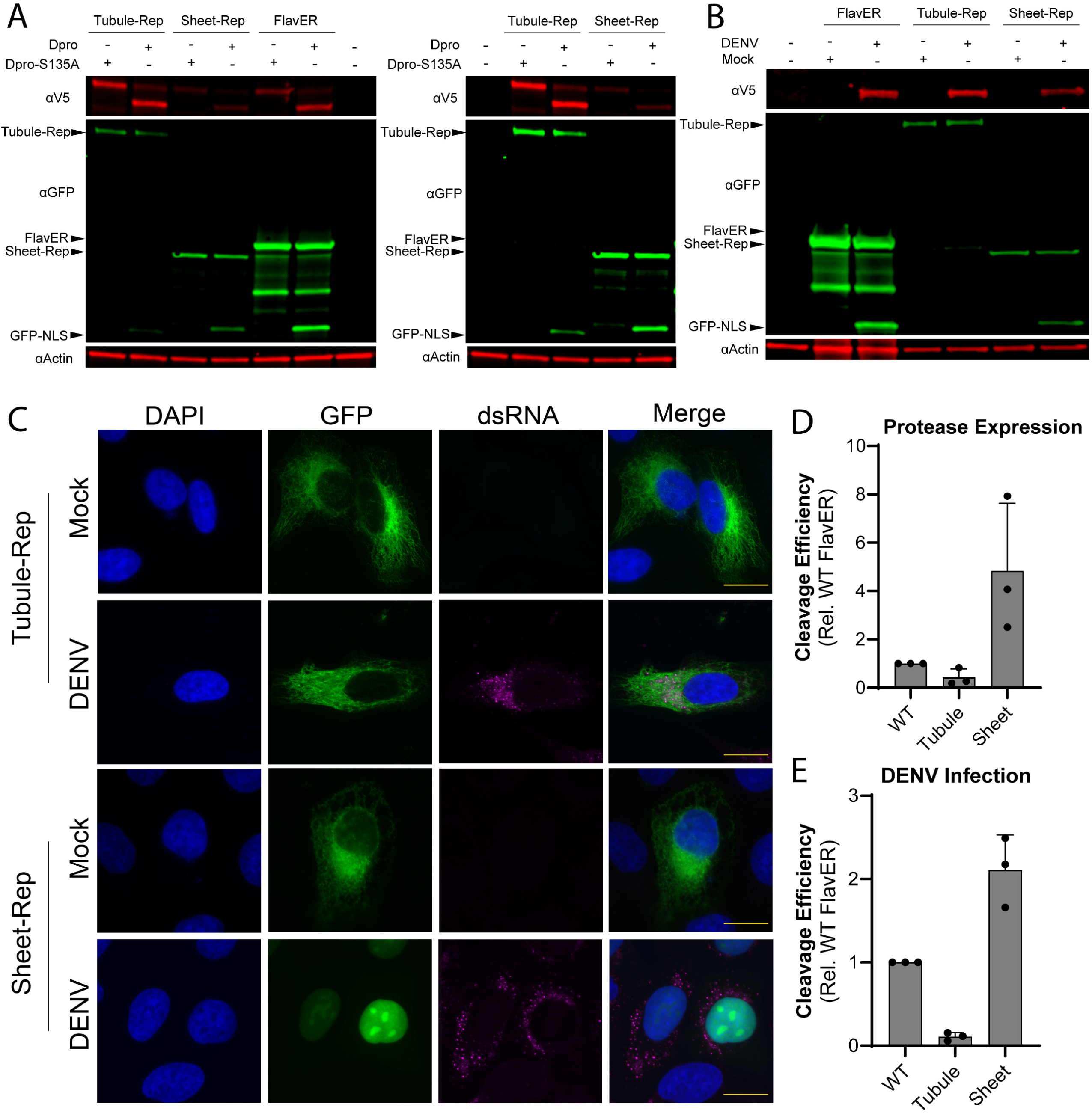
Substrate localization to ER subdomains as a molecular determinant for cleavage. A.) *Left*, immunoblot for V5, GFP, and actin of U2OS cells co-transfected with the indicated reporter construct and DENV protease. Labeled arrows denote the full-length (Tubule-Rep, FlavER, Sheet-Rep) and cleaved (GFP-NLS) reporter construct. *Right*, cropped immunoblot with increased brightness to emphasize cleavage band (GFP-NLS) of Tubule-Rep. B.) Immunoblot for DENV NS3, GFP, and actin of U2OS cells co-transfected with the indicated reporter construct and mock or DENV-infected. Labeled arrows denote the full-length (Tubule-Rep, FlavER, Sheet-Rep) and cleaved (GFP-NLS) reporter construct. C.) IF microscopy of mock or DENV-infected U2OS cells expressing GFP-Tubule-Rep (top two panels) or GFP-Sheet-Rep (bottom two panels). Reporter constructs shown in green, dsRNA shown in magenta, and nuclei are stained with DAPI and shown in blue. Scale bars represent 20 um. D-E.) Quantification of cleavage assays from reporter-expressing cells transfected with Dpro (D) or infected with DENV at an MOI of 3 (E). Data are presented as efficiency of cleavage determined by intensity ratios in arbitrary units (AU) of the cleaved to total reporter signal relative to viral protease expression. Quantification represented as cleavage efficiency relative to the FlavER-WT construct. Each bar represents the average cleavage efficiency of three experiments per condition and black circles indicate the cleavage efficiency calculated for each replicate.

### Orthoflavivirus proteases exhibit sequence-specific cleavage efficiency profiles

We next sought to explore the primary sequence of the substrate cut site as a molecular determinant of cleavage by orthoflavivirus proteases. It is known that the majority of the cytoplasmic sequences of the viral polyprotein cleavage motifs include two basic residues followed by a small amino acid (30, 32, 60, 61). However, when looking at the sequences present at the different cleavage junctions within the polyproteins of DENV, ZIKV, WNV, and YFV, there is a notable degree of variance in the motifs that are processed at different junctions within a single polyprotein, as well as differences in the sequences cleaved at the same junction across different viral species (Fig. 5A). To assess the degree of variability in the cleavage efficiency of the protease recognition motifs located in the cytoplasmic junctions of the flavivirus polyproteins, we designed 8 reporter constructs encoding the non-repetitive cleavage sequences within the cytoplasmic junctions of the DENV, ZIKV, WNV, and YFV polyproteins (Fig. 5A, B). We then performed cleavage assays where one of the eight reporter constructs was co-expressed with a DENV, ZIKV, WNV, or YFV protease. Quantification of the immunoblots from these experiments revealed that each flavivirus protease has distinct cleavage efficiencies for the different sequences present within their polyprotein junctions and that these preferences varied across viral species (Fig. 5C-F). For example, the ZIKV and WNV proteases had very similar graph patterns, while the DENV and WNV graph patterns were nearly opposite. However, the most intriguing detail from this data is that the sequence within each viral polyprotein that was processed least efficiently by its protease was located at the same cytoplasmic junction: between NS4A and the 2K peptide (Fig. 5B-F). This suggests that there is an essential role for the suboptimal cleavage of the motif at this position within the viral polyprotein. Together, these results revealed that each flavivirus protease has a unique cleavage profile and suggests an evolutionary advantage to maintaining sequences of varying cleavage efficiencies at certain polyprotein junctions.

**Figure 5.**
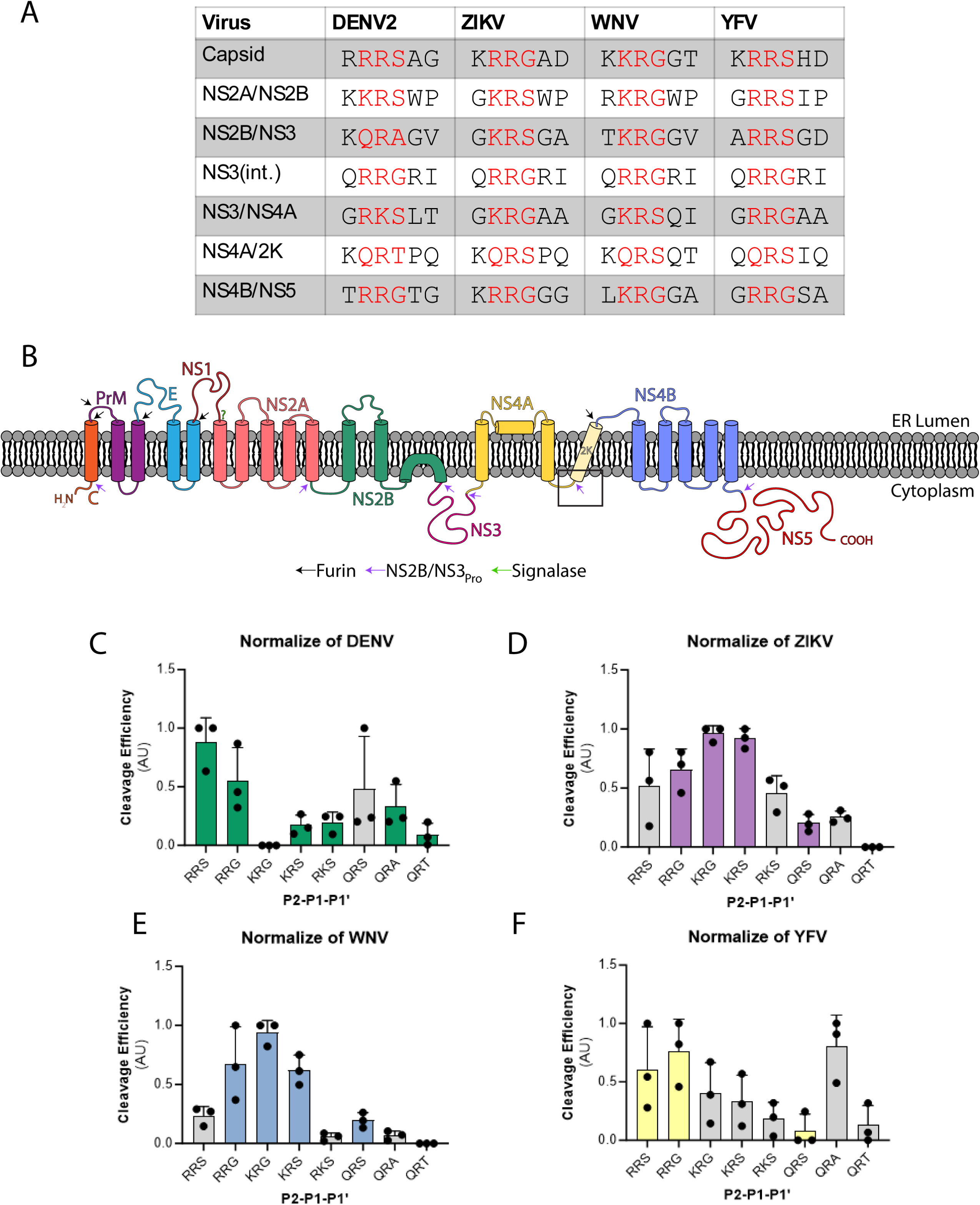
Sequence-specific cleavage efficiency profiles. A.) Model of a pre-processed flavivirus polyprotein translated into the membrane of the ER. Purple arrows represent cytoplasmic junctions cleaved by the viral protease (NS2B/NS3pro). Green arrows indicate luminal junctions cleaved by host signalase. Black arrow represents the luminal junction cleaved by host furin. Cleavage junction between NS4A and the 2K peptide is marked by the black box. B.) Table representing the cleavage motifs at the seven cytoplasmic cleavage junctions processed by the viral proteases in the DENV2, ZIKV, WNV, and YFV polyproteins. Solved three amino acid cleavage motifs indicated in red letters. C-F.) Cleavage efficiency plots as determined by immunoblot for eight indicated P2-P1’ sequences present in the cytoplasmic junctions of DENV (C), ZIKV (D), WNV (E), and YFV (F) polyproteins. Bars in color indicate the cleavage junction sequences present within the polyproteins that encode the tested protease. Gray bars represent sequences absent from the polyproteins of the indicated protease. Data is presented as efficiency of cleavage determined by intensity ratios in arbitrary units (AU) of the cleaved to total reporter signal relative to viral protease expression. Each bar represents the average cleavage efficiency of three experiments per condition and black circles indicate the cleavage efficiency calculated for each replicate.

### Cleavage of the NS4A|2K sequence is significantly delayed

The cleavage profiles provide insight into the viral proteases’ cleavage efficiency of different substrates after a 24-hour post-transfection timepoint. However, we hypothesized that the timing of cleavage would be the most important factor in the context of polyprotein processing. Therefore, a more detailed way to study viral protease cleavage efficiency is to evaluate the cleavage kinetics of our reporters during infection. To investigate the intracellular cleavage kinetics, we can adapt our reporter system to long-term, time-lapse imaging of living cells (41). The visual advantage of our reporter construct is the nuclear translocation of the fluorescent reporter signal that occurs in response to cleavage of the protease recognition motif. Using this reporter system in combination with live-cell imaging allows us to study the efficiency of intracellular reporter cleavage over time by measuring the increase in reporter signal in the nucleus of the cell throughout viral infection (41).

In order to directly compare the intracellular cleavage kinetics of varying sequences, we can design reporters encoding different fluorescent proteins (GFP and mCherry) fused upstream of the two cleavage motifs. The fluorescent protein fused downstream of the TM domain in the original FlavER design was replaced with a NanoLuciferase to ensure that each reporter expresses a single fluorescent protein and that proper topology and localization of the reporter constructs are maintained. This method allowed us to visualize and quantify the nuclear translocation of the two fluorescent signals that occurs as a result of proteolytic cleavage of the respective reporter sequences within a single infected cell co-expressing the reporters. To utilize this assay, we first validated its efficacy by confirming that there was no inherent difference in the rate of nuclear translocation between GFP and mCherry. To confirm this, we designed two reporters encoding the same protease recognition sequence fused to different fluorescent proteins (GFP-QR|T, mCh-QR|T) and expressed them in cells together. The co-expressing cells were then infected with DENV 4 hours prior to live-cell imaging for a 20-hour timespan, where images were captured in 20-minute intervals and compiled into movies. Using this method, we began to observe nuclear translocation of both GFP and mCherry at about 12 HPI, and the nuclear signal steadily increased as infection progressed. (Fig. 6A, Movie 1). Quantification of the signal intensities of both GFP and mCherry in the nucleus in each captured frame revealed no difference in the rate of nuclear translocation between the two fluorescent signals (Fig. 6B). We further calculated the average time at which the 50% maximum signal was reached in the nucleus (HPI50) during infection and observed no significant difference between GFP and mCherry (Fig. 6C). Together, these results indicate that this method can be used to compare the intracellular cleavage kinetics of two cleavage motifs fused to the different fluorescent proteins.

**Figure 6.**
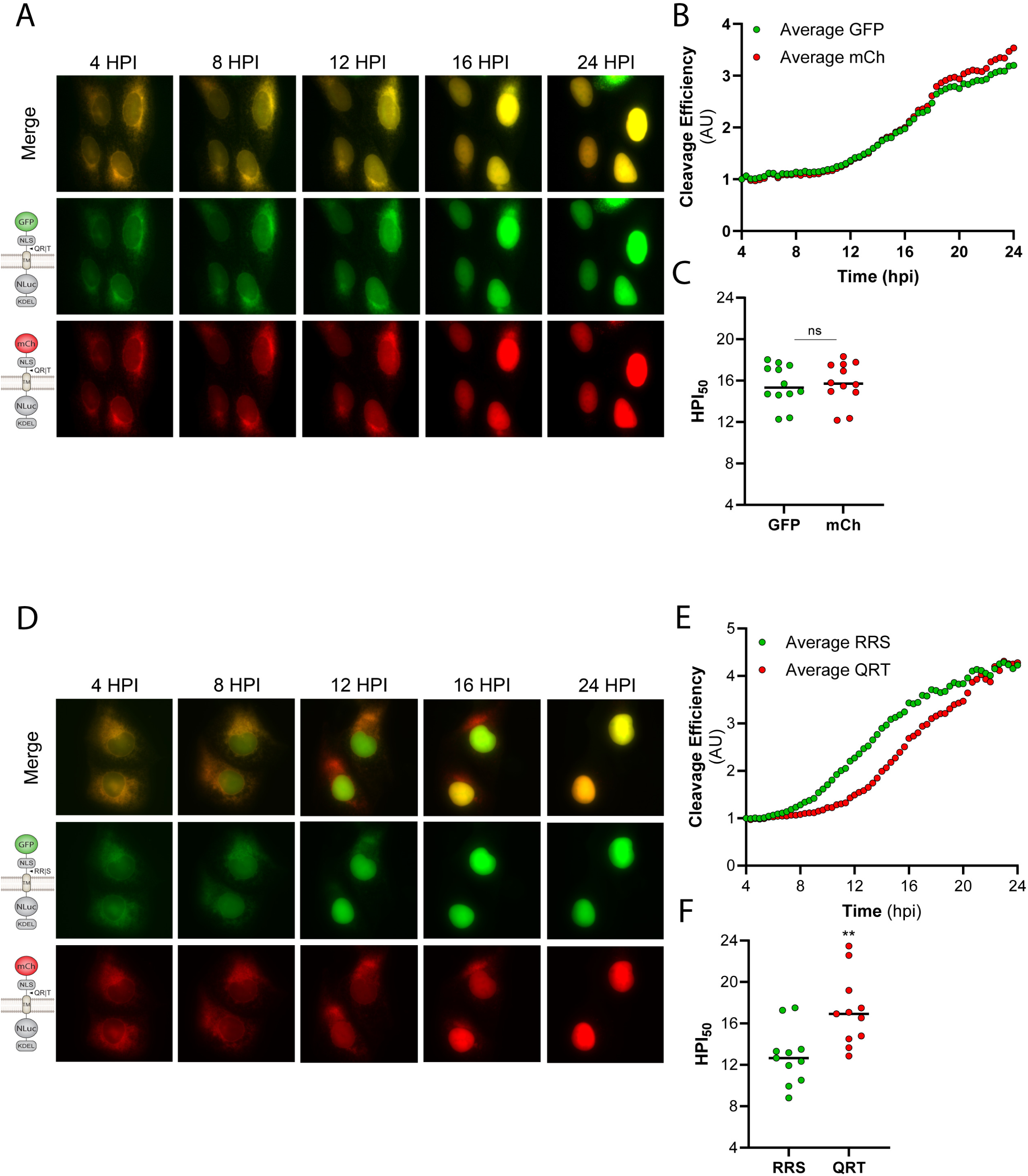
Comparative cleavage kinetics of capsid versus NS4A|2K sequences. A,D.) Representative time-points of image series from live cell imaging of DENV-infected cells co-expressing (A) GFP-QR|T or (D) GFP-RR|S (middle) and mCh-QR|T (bottom) reporters. Merged images in top panels. B,E.) Quantification of cleavage efficiencies of (B) GFP-QRT or (E) GFP-RRS vs. mCh-QRT reporters, defined as the reporter signal intensity in the nucleus of infected cells co-expressing the two constructs (n=12). Individual data points represent the average reporter intensity in the nucleus at the indicated time point. C,F.) Graphs representing the times at which the nucleus reaches the 50% maximum intensity (HPI50) of each reporter signal as determined by nonlinear regression analysis for each DENV-infected cell. Values for GFP versus mCherry reporters compared using unpaired t-tests.

To further understand the significance of the variable cleavage sequences present in flavivirus polyproteins, we aimed to directly compare the cleavage kinetics between the two protease recognition motifs found to be the most and least efficiently processed by the DENV protease. As shown in Figure 5C, the DENV protease processed the RR|S sequence most efficiently and the QR|T sequence least efficiently. These motifs are located at the cytoplasmic capsid junction and NS4A|2K junction of the viral polyprotein, respectively. Therefore, we designed a reporter encoding a GFP upstream of the RR|S recognition motif (GFP-RR|S) to co-express with the mCh-QR|T reporter in cells to be infected with DENV (Fig. 6D). Using the same technique described previously, we found a distinct difference in the timing of the onset of nuclear translocation between the GFP and the mCherry signals (Movie 2). On average, we found that GFP began translocating at approximately 8 HPI, while mCherry did not begin translocating until approximately 12 HPI. Further, we observed complete nuclear translocation of GFP by ∼16 HPI and of mCherry by the endpoint of ∼24 HPI, on average (Fig. 6D,E). Calculation of HPI50 during infection resulted in ∼13 HPI for GFP and ∼17 HPI for mCherry (Fig. 6F). Together, these results suggest a significant, approximately 4-hour, delay in DENV protease cleavage of the QR|T sequence, located at the NS4A|2K junction within the viral polyprotein, compared to the RR|S motif.

### QR|T cleavage efficiency increases in later DENV infection

Next, we wanted to validate our cleavage kinetics results with our standard cleavage assays to highlight the importance of timing in the cleavage of intracellular sequences, particularly when considering polyprotein processing in the context of infection. To corroborate the live-cell imaging results, we independently expressed GFP reporters encoding either the QR|T or RR|S motifs in U2OS cells. These cells were then infected with DENV at an MOI of 3, and lysates were collected at 12 and 24 HPI. We observed that in our early infection time point, there is far more efficient cleavage of our RR|S reporter compared to the QR|T reporter, as indicated by the more robust cleavage band shown by immunoblot. However, by 24 HPI, we observed a strong cleavage band of our QR|T reporter that nearly mirrored the RR|S reporter cleavage efficiency (Fig. 7A). Signal intensity quantification of the intracellular cleavage assays paralleled these results. We observed a significant defect in QR|T reporter cleavage compared to that of the RR|S reporter motif at 12 HPI, yet there was no significant difference in cleavage efficiency between the two reporters by 24 HPI (Fig. 7B). This data accurately corresponds with the trend observed in our live-cell cleavage kinetics assay using DENV infection and emphasizes the impact that progression of infection over time has on viral protease activity.

**Figure 7.**
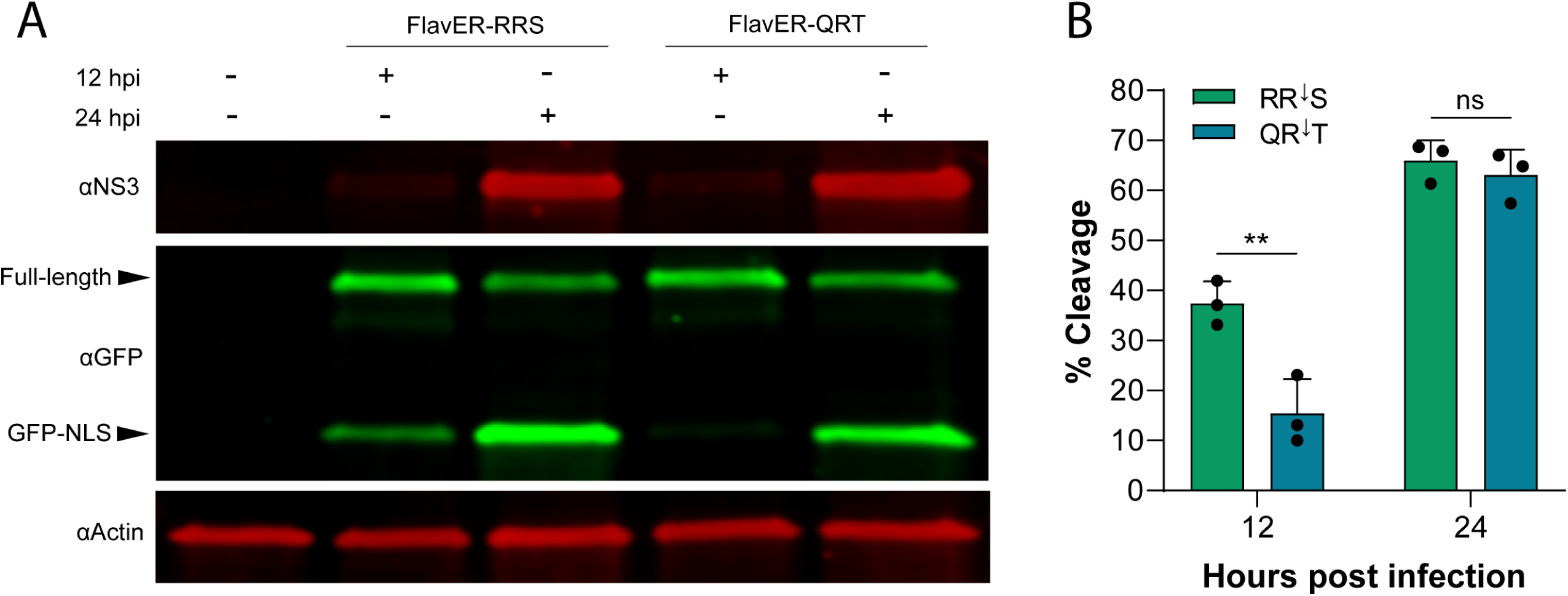
Cleavage of capsid versus NS4A|2K sequences at 12 and 24 hours post infection. A.) Immunoblot for NS3, GFP, and actin of DENV-infected U2OS cells expressing GFP-RRS or GFP-QRT. Cells were infected with DENV at an MOI of 3 and lysed at either 12 or 24 hours post infection. Labeled arrows denote the full-length and cleaved (GFP-NLS) reporter constructs. B.) Quantification of immunoblot defined as % cleavage of the reporter at 12 and 24 hours post infection. Statistical significance determined by one-way ANOVA.

### Altering cleavage kinetics at the NS4A|2K junction is lethal

We next wanted to understand the effect of polyprotein cleavage kinetics on the infectivity of flaviviruses. To investigate this process, we employed our recently developed DNA-launched DENV infectious clone that allows for the recovery of high viral titers after transfection into cells (40). We replaced the WT (QR|T) residues between NS4A and the 2K peptide with the cleavage motifs found to be the most efficiently processed by the DENV protease (RR|S) to increase the rate of cleavage at this junction within the viral polyprotein (Fig. 5C, Fig. 8A). We observed efficient recovery of WT virus, but we were unable to detect any foci from the cells incubated with the mutant supernatants. To see if this effect was conserved in another orthoflavivirus species, we utilized a ZIKV infectious clone (39). Using the same method described previously, we found that increasing the rate of cleavage at the NS4A|2K junction within the ZIKV polyprotein (QR|S → KR|G) also had lethal effects on viral recovery. These results suggest that suboptimal cleavage kinetics between NS4A and the 2K peptide within flavivirus polyproteins is essential for viral fitness.

**Figure 8.**
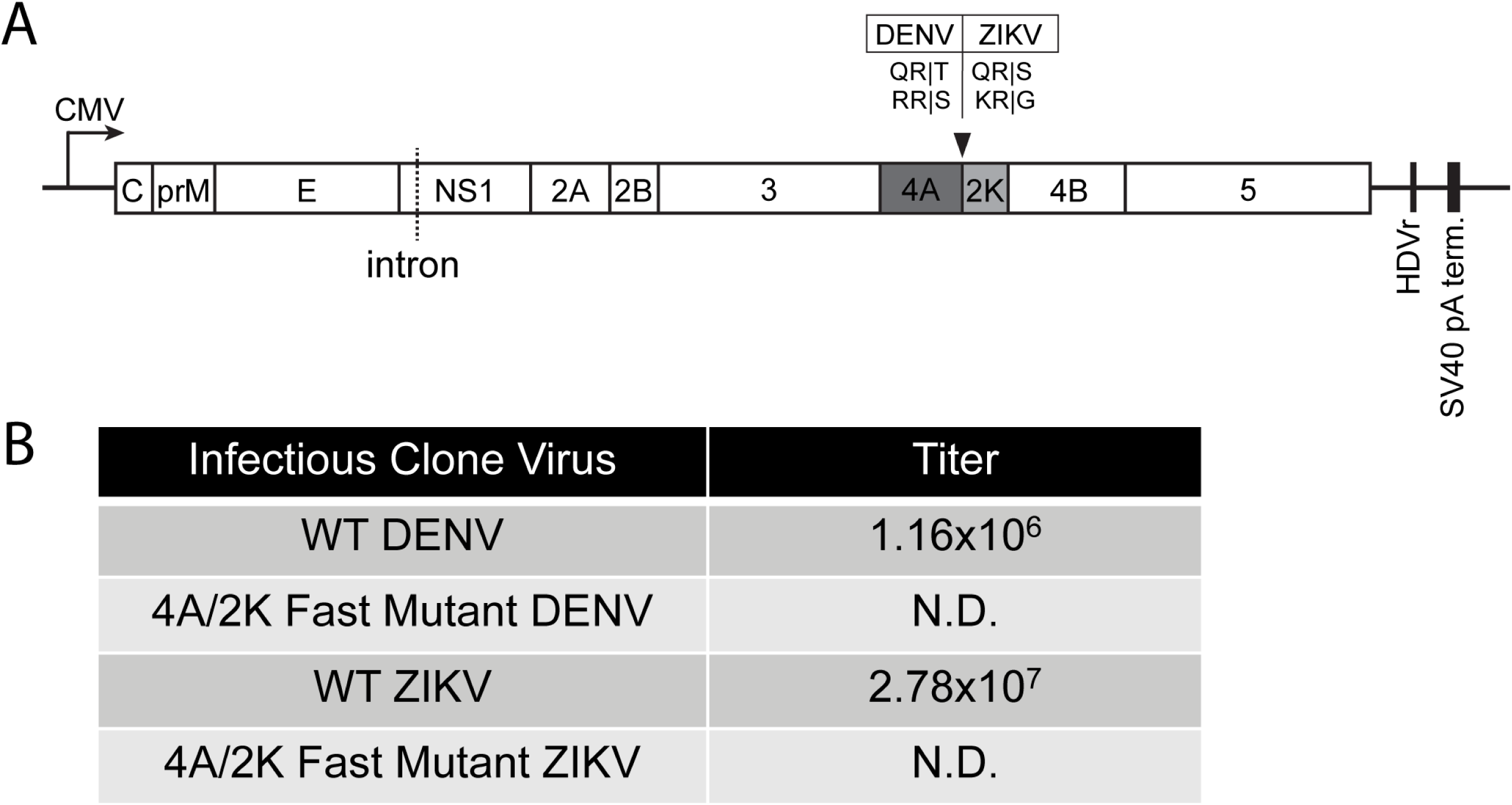
Altering cleavage kinetics at the NS4A|2K junction is lethal for DENV and ZIKV. A.) Linear map of DNA-launched DENV and ZIKV infectious clones. Viral genomes lie between a CMV promotor and an HDV ribozyme (HDVr). An artificial intron is introduced in the NS1 gene segment to promote stability in bacteria. NS4A and 2K gene segments represented as dark and light gray boxes, respectively. Black arrow indicates the cleavage junction where mutations were introduced. B.) Rescue of recombinant virus encoding the WT or fast cleavage sequence at the NS4A|2K junction within the polyproteins of the DENV or ZIKV infectious clones. N.D. = Not detectable.

## Discussion

Orthoflavivirus proteases are essential protein complexes responsible for processing the viral polyprotein into its individual subunits that perform the functions to mediate infection. These proteases have also been shown to interact with different host proteins in order to establish a favorable environment for replication (62, 63). Because productive infections cannot be established without proper function of the viral protease, this protein complex has long been an attractive target for the development of antiviral therapeutics. However, no orthoflavivirus protease inhibitors have been successful in the clinic; thus, a further understanding of its molecular functions is needed for the development of targeted antivirals (37). In this study, we sought to identify the subcellular determinants involved in flavivirus protease-induced cleavage of substrates. We first used our tractable protease activity reporter system to identify two previously uncharacterized molecular factors associated with substrate cleavage by the DENV protease: membrane proximity of the recognition motif and ER subdomain localization of the substrate. We then investigated primary sequence specificity for multiple flavivirus proteases as this was the only previously characterized cleavage determinant prior to this study. We found that although each flavivirus protease had a unique cleavage profile, they all shared the characteristic of processing the sequence located at the NS4A|2K junction with the poorest efficiency. This led us to uncover the importance of the rate of this cleavage event for viral infectivity. Overall, this report investigated the molecular determinants for flavivirus protease cleavage and further revealed that cleavage efficiency of the viral polyprotein plays a pivotal role in viral fitness.

Our FlavER reporter system is a valuable tool for studying intracellular cleavage specificity of flavivirus proteases. Here, we show the reporter platform can accommodate various modifications to allow for the detailed investigation of substrate cleavage efficiency by viral proteases. First, we found that increasing the distance of the cut site from the TM domain in our reporter led to a correlative decrease in cleavage efficiency by the DENV protease (Fig. 2). This molecular determinant is consistent with what is proposed about the cytoplasmic cleavage junctions within the viral polyprotein. The majority of the cytoplasmic polyprotein cleavage junctions are assumed to be within less than 10 amino acids from the ER membrane (Fig. 2B) (64–67). The junctions presumed to be farther from the ER membrane, NS3int and NS3-NS4A, have both been shown to be strictly intramolecular cleavage events which could account for the tolerability of greater membrane distances (68). Further, several studies have revealed that cleavage at both the internal NS3 site as well as the NS3|NS4A junction are less efficient and kinetically slower than other junctions within the polyprotein which could be attributed to the membrane proximity of the cut sites (69–71).

We also found that the localization to specific subdomains of the ER had profound effects on the cleavage efficiency of the reporter constructs. We observed that cleavage of our ER-sheet reporter was far more efficient than cleavage of the ER-tubule reporter during both DENV infection and protease overexpression (Fig. 4). Notably, we observed that during DENV infection, the viral protease is unable to cleave the reporter localized to ER tubules (Fig. 4B). This observation is consistent with reports that showed dsRNA, viral proteins, and assembled flavivirus particles localized specifically in dilated ER cisternae (72, 73). These results reveal the impact of the distinct localization of DENV protease activity within the ER while emphasizing the importance of validating protease expression assays with infection studies.

The final molecular determinant we investigated further was the primary sequence of the viral protease recognition motif. Many studies have reported on the cleavage specificity of flavivirus proteases using purified recombinant proteases and *in vitro* enzymatic assays (33, 74–77). Though these are valuable studies for gaining an understanding of protease activity, these methods take the viral protease out of the context of the cell and away from the membrane that it typically associates with. This limitation is eliminated with the use of our reporter system as we test the cleavage of intracellular substrates with a more native form of the viral protease that is exposed to natural cellular constraints. Using our reporter platform, we identified that the DENV, ZIKV, WNV, and YFV proteases each had distinct cleavage specificity patterns (Fig. 5). Understanding these cleavage profiles has potential to provide insight on how cleavage efficiency at certain cytoplasmic junctions affects the coordinated processing of the viral polyprotein. A few studies have determined that certain polyprotein cleavage events are prerequisites for processing at another junction to occur; however, the chronological sequence of cleavage events of the flavivirus polyproteins is largely unknown and warrants further investigation (78–80).

These results are also valuable for establishing parameters to use when evaluating potential host protein candidates that may be targeted for cleavage by orthoflavivirus proteases. Previous studies have uncovered three ER-resident host proteins known to be cleaved by multiple flavivirus proteases: stimulator of interferon genes (STING), reticulophagy receptor 1 (RETREG1), and diacylglycerol O-acyltransferase 2 (DGAT2). Both STING and RETREG1 are known to restrict flavivirus infection (42, 81–83). In response, multiple flavivirus proteases evolved the ability to cleave these host proteins at non-consensus cleavage sequences in order to inhibit their antiviral functions (42, 84–86). DGAT2 is unique from STING and RETREG1 in that cleavage by the viral protease stabilizes this host protein for the virus to hijack its function as a proviral factor (87). DGAT2 is known to promote the formation of lipid droplets which have been shown to serve as an energy source of viral replication (88–90). Thus, enhancing the stability of DGAT2 allows for lipid droplet accumulation that creates a favorable environment for viral replication. Other non-ER-resident host proteins, specifically factors localized to the mitochondria, have also been shown to be cleaved by flavivirus proteases (91, 92). Our results would suggest that these proteins are likely interacting with other membrane proximal host or viral proteins that allow the protease to access the cleavage site. Identifying host proteins that are cleaved by flavivirus proteases is a promising area of research as it provides an understanding of how this viral protein complex interacts with cellular factors in order to promote replication. The results of this study can support these research efforts by identifying important parameters to acknowledge when evaluating cleavage targets. Considering the ER subdomain localization of the host protein and the primary sequence and membrane proximity of the potential cleavage site could allow for a more accurate assessment of its cleavage probability.

When investigating primary sequence specificity, we also identified that each protease processed the sequence located at the NS4A|2K junction least efficiently and that proteolytic processing of the NS4A|2K (QR|T) sequence was significantly delayed compared to the capsid cleavage sequence (RR|S) during DENV infection (Fig. 5,6D-F). Notably, we observed that in contrast to the cleavage assays with exogenous protease expression, the degree of QR|T cleavage nearly matches that of RR|S at 24 hours in the context of DENV infection (Fig. 7). These results indicate that cleavage of the NS4A|2K sequence may require a specific environment consisting of a high concentration of protease and/or infection-induced membrane rearrangements, which cannot be simulated in protease transfections. Interestingly, ultrastructural studies of infected cells have indicated these viruses induce temporal changes to the ER membrane that begin with the formation of membrane packets that go on to enclose de novo vesicles as infection progresses (93). Thus, it is possible that efficiency-regulated cleavage of the viral polyprotein is linked to transitional virus-induced membrane alterations.

Lastly, we also sought to determine if the delayed cleavage event at the NS4A|2K junction of the flavivirus polyprotein was critical for viral fitness and found that introducing a high efficiency cleavage site between NS4A and the 2K peptide led to complete loss of infectious viral progeny (Fig. 8). These results suggest that there is an essential purpose for the delay in proteolytic processing at this particular junction. One possibility for this function is that a cleavage delay is required for efficient replication to occur during early infection. This hypothesis is supported by recent studies that have shown that the unprocessed NS4A-2K-NS4B precursor operates as a functional intermediate that interacts with other viral proteins. Specifically, two separate reports found that the NS4A-2K-NS4B intermediate interacts with both NS1 and NS3 and that these interactions are required for DENV replication (94–96). These studies bolster the idea that the purpose behind the delay in NS4A|2K processing is due to the functions performed by NS4A-2K-NS4B in its pre-cleaved form. This could suggest that premature separation of NS4A and NS4B due to the introduction of a more efficient cleavage sequence may interrupt these crucial interactions and inhibit replication resulting in the loss of fitness (Fig. 7). Future studies are targeted at further understanding the impact of temporal polyprotein processing on virus infection.

Collectively, this report identified membrane proximity and ER subdomain localization as molecular determinants associated with substrate cleavage by orthoflavivirus proteases and highlighted the impact of sequence specificity on polyprotein processing and subsequent viral fitness. Our data further provides a set of parameters to consider when evaluating promising host protein candidates that may be targeted by these viral proteases. Together, these results advance our understanding of the intracellular molecular activities of orthoflavivirus proteases which can contribute to closing the gap in knowledge of their complex mechanisms to support the development of targeted antiviral therapeutics.

## Acknowledgements

This research was supported by National Institutes of Health F31-AI183737 (L.C.) and R35-GM150638 (N.J.L.), as well as the University of Alabama at Birmingham Heersink School of Medicine (N.J.L. and C.M.P.). We thank Carolyne Coyne (Duke University) and Kevin Harrod (University of Alabama at Birmingham) for reagents.

## References

1. Souza-Neto JA, Powell JR, Bonizzoni M. 2019. Aedes aegypti vector competence studies: A review. Infection, Genetics and Evolution 67:191–209.

2. Valderrama A, Díaz Y, López-Vergès S. 2017. Interaction of Flavivirus with their mosquito vectors and their impact on the human health in the Americas. Biochem Biophys Res Commun 492:541–547.

3. Weaver SC, Charlier C, Vasilakis N, Lecuit M. 2018. Zika, Chikungunya, and Other Emerging Vector-Borne Viral Diseases. Annu Rev Med 69:395–408.

4. Spadar A, Phelan JE, Benavente ED, Campos M, Gomez LF, Mohareb F, Clark TG, Campino S. 2021. Flavivirus integrations in Aedes aegypti are limited and highly conserved across samples from different geographic regions unlike integrations in Aedes albopictus. Parasit Vectors 14:332.

5. Hale GL. 2023. Flaviviruses and the Traveler: Around the World and to Your Stage. A Review of West Nile, Yellow Fever, Dengue, and Zika Viruses for the Practicing Pathologist. Modern Pathology 36:100188.

6. Liang Y, Dai X. 2024. The global incidence and trends of three common flavivirus infections (Dengue, yellow fever, and Zika) from 2011 to 2021. Front Microbiol 15.

7. Childs ML, Lyberger K, Harris M, Burke M, Mordecai EA. 2024. Climate warming is expanding dengue burden in the Americas and Asia. medRxiv 10.1101/2024.01.08.24301015.

8. Bouri N, Sell TK, Franco C, Adalja AA, Henderson DA, Hynes NA. 2012. Return of Epidemic Dengue in the United States: Implications for the Public Health Practitioner. Public Health Reports® 127:259–266.

9. Butterworth MK, Morin CW, Comrie AC. 2017. An Analysis of the Potential Impact of Climate Change on Dengue Transmission in the Southeastern United States. Environ Health Perspect 125:579–585.

10. Pierson TC, Diamond MS. 2020. The continued threat of emerging flaviviruses. Nat Microbiol 5:796–812.

11. Simo FBN, Bigna JJ, Kenmoe S, Ndangang MS, Temfack E, Moundipa PF, Demanou M. 2019. Dengue virus infection in people residing in Africa: a systematic review and meta-analysis of prevalence studies. Sci Rep 9:13626.

12. Uno N, Ross TM. 2018. Dengue virus and the host innate immune response. Emerg Microbes Infect 7:1–11.

13. Rossi SL, Ross TM, Evans JD. 2010. West Nile Virus. Clin Lab Med 30:47–65.

14. Duarte G, Moron AF, Timerman A, Fernandes CE, Mariani Neto C, Almeida Filho GL de, Werner Junior H, Espírito Santo HFB do, Steibel JAP, Bortoletti Filho J, Andrade JBB de, Burlá M, Silva de Sá MF, Busso NE, Giraldo PC, Moreira de Sá RA, Passini Junior R, Mattar R, Francisco RPV. 2017. Zika Virus Infection in Pregnant Women and Microcephaly. Revista Brasileira de Ginecologia e Obstetrícia / RBGO Gynecology and Obstetrics 39:235–248.

15. Guo C, Zhou Z, Wen Z, Liu Y, Zeng C, Xiao D, Ou M, Han Y, Huang S, Liu D, Ye X, Zou X, Wu J, Wang H, Zeng EY, Jing C, Yang G. 2017. Global Epidemiology of Dengue Outbreaks in 1990–2015: A Systematic Review and Meta-Analysis. Front Cell Infect Microbiol 7.

16. Daep CA, Muñoz-Jordán JL, Eugenin EA. 2014. Flaviviruses, an expanding threat in public health: focus on dengue, West Nile, and Japanese encephalitis virus. J Neurovirol 20:539–560.

17. Hameed M, Wahaab A, Shan T, Wang X, Khan S, Di D, Xiqian L, Zhang J-J, Anwar MN, Nawaz M, Li B, Liu K, Shao D, Qiu Y, Wei J, Ma Z. 2024. Corrigendum: A metagenomic analysis of mosquito virome collected from different animal farms at Yunnan–Myanmar border of China. Front Microbiol 15.

18. Thomas SJ, Yoon I-K. 2019. A review of Dengvaxia®: development to deployment. Hum Vaccin Immunother 15:2295–2314.

19. Ishikawa T, Yamanaka A, Konishi E. 2014. A review of successful flavivirus vaccines and the problems with those flaviviruses for which vaccines are not yet available. Vaccine 32:1326–1337.

20. Heinz FX, Stiasny K. 2012. Flaviviruses and flavivirus vaccines. Vaccine 30:4301–4306.

21. Ngo AM, Shurtleff MJ, Popova KD, Kulsuptrakul J, Weissman JS, Puschnik AS. 2019. The ER membrane protein complex is required to ensure correct topology and stable expression of flavivirus polyproteins. Elife 8.

22. Mazeaud C, Freppel W, Chatel-Chaix L. 2018. The Multiples Fates of the Flavivirus RNA Genome During Pathogenesis. Front Genet 9.

23. Amberg SM, Rice CM. 1999. Mutagenesis of the NS2B-NS3-Mediated Cleavage Site in the Flavivirus Capsid Protein Demonstrates a Requirement for Coordinated Processing. J Virol 73:8083–8094.

24. Rice CM. 1996. Flaviviridae: the viruses and their replication. 1:931–960.

25. Yusof R, Clum S, Wetzel M, Murthy HMK, Padmanabhan R. 2000. Purified NS2B/NS3 Serine Protease of Dengue Virus Type 2 Exhibits Cofactor NS2B Dependence for Cleavage of Substrates with Dibasic Amino Acids in Vitro. Journal of Biological Chemistry 275:9963–9969.

26. Wahaab A, Liu K, Hameed M, Anwar MN, Kang L, Li C, Ma X, Wajid A, Yang Y, Khan UH, Wei J, Li B, Shao D, Qiu Y, Ma Z. 2021. Identification of Cleavage Sites Proteolytically Processed by NS2B-NS3 Protease in Polyprotein of Japanese Encephalitis Virus. Pathogens 10:102.

27. Amberg SM, Rice CM. 1999. Mutagenesis of the NS2B-NS3-Mediated Cleavage Site in the Flavivirus Capsid Protein Demonstrates a Requirement for Coordinated Processing. J Virol 73:8083–8094.

28. Xing H, Xu S, Jia F, Yang Y, Xu C, Qin C, Shi L. 2020. Zika NS2B is a crucial factor recruiting NS3 to the ER and activating its protease activity. Virus Res 275:197793.

29. Wu C-F, Wang S-H, Sun C-M, Hu S-T, Syu W-J. 2003. Activation of dengue protease autocleavage at the NS2B–NS3 junction by recombinant NS3 and GST–NS2B fusion proteins. J Virol Methods 114:45–54.

30. Falgout B, Pethel M, Zhang YM, Lai CJ. 1991. Both nonstructural proteins NS2B and NS3 are required for the proteolytic processing of dengue virus nonstructural proteins. J Virol 65:2467–2475.

31. Chambers TJ, Nestorowicz A, Amberg SM, Rice CM. 1993. Mutagenesis of the yellow fever virus NS2B protein: effects on proteolytic processing, NS2B-NS3 complex formation, and viral replication. J Virol 67:6797–6807.

32. Chambers TJ, Weir RC, Grakoui A, McCourt DW, Bazan JF, Fletterick RJ, Rice CM. 1990. Evidence that the N-terminal domain of nonstructural protein NS3 from yellow fever virus is a serine protease responsible for site-specific cleavages in the viral polyprotein. Proceedings of the National Academy of Sciences 87:8898–8902.

33. Shiryaev SA, Kozlov IA, Ratnikov BI, Smith JW, Lebl M, Strongin AY. 2007. Cleavage preference distinguishes the two-component NS2B–NS3 serine proteinases of Dengue and West Nile viruses. Biochemical Journal 401.

34. Wahaab A, Mustafa BE, Hameed M, Stevenson NJ, Anwar MN, Liu K, Wei J, Qiu Y, Ma Z. 2021. Potential Role of Flavivirus NS2B-NS3 Proteases in Viral Pathogenesis and Anti-flavivirus Drug Discovery Employing Animal Cells and Models: A Review. Viruses 14:44.

35. Luo D, Vasudevan SG, Lescar J. 2015. The flavivirus NS2B–NS3 protease–helicase as a target for antiviral drug development. Antiviral Res 118:148–158.

36. Diamond MS, Pierson TC. 2015. Molecular Insight into Dengue Virus Pathogenesis and Its Implications for Disease Control. Cell 162:488–92.

37. Cavina L, Bouma MJ, Gironés D, Feiters MC. 2024. Orthoflaviviral Inhibitors in Clinical Trials, Preclinical In Vivo Efficacy Targeting NS2B-NS3 and Cellular Antiviral Activity via Competitive Protease Inhibition. Molecules 29:4047.

38. Samrat SK, Xu J, Li Z, Zhou J, Li H. 2022. Antiviral Agents against Flavivirus Protease: Prospect and Future Direction. Pathogens 11:293.

39. Schwarz MC, Sourisseau M, Espino MM, Gray ES, Chambers MT, Tortorella D, Evans MJ. 2016. Rescue of the 1947 Zika Virus Prototype Strain with a Cytomegalovirus Promoter-Driven cDNA Clone. mSphere 1.

40. Holliday M, Corliss L, Lennemann NJ. 2023. Construction and Rescue of a DNA-Launched DENV2 Infectious Clone. Viruses 15:275.

41. Corliss L, Holliday M, Lennemann NJ. 2022. Dual-fluorescent reporter for live-cell imaging of the ER during DENV infection. Front Cell Infect Microbiol 12.

42. Lennemann NJ, Coyne CB. 2017. Dengue and Zika viruses subvert reticulophagy by NS2B3-mediated cleavage of FAM134B. Autophagy 13.

43. Abramson J, Adler J, Dunger J, Evans R, Green T, Pritzel A, Ronneberger O, Willmore L, Ballard AJ, Bambrick J, Bodenstein SW, Evans DA, Hung C-C, O’Neill M, Reiman D, Tunyasuvunakool K, Wu Z, Žemgulytė A, Arvaniti E, Beattie C, Bertolli O, Bridgland A, Cherepanov A, Congreve M, Cowen-Rivers AI, Cowie A, Figurnov M, Fuchs FB, Gladman H, Jain R, Khan YA, Low CMR, Perlin K, Potapenko A, Savy P, Singh S, Stecula A, Thillaisundaram A, Tong C, Yakneen S, Zhong ED, Zielinski M, Žídek A, Bapst V, Kohli P, Jaderberg M, Hassabis D, Jumper JM. 2024. Accurate structure prediction of biomolecular interactions with AlphaFold 3. Nature 630:493–500.

44. Humphrey W, Dalke A, Schulten K. 1996. VMD: visual molecular dynamics. J Mol Graph 14:33–8, 27–8.

45. Phillips JC, Hardy DJ, Maia JDC, Stone JE, Ribeiro J V, Bernardi RC, Buch R, Fiorin G, Hénin J, Jiang W, McGreevy R, Melo MCR, Radak BK, Skeel RD, Singharoy A, Wang Y, Roux B, Aksimentiev A, Luthey-Schulten Z, Kalé L V, Schulten K, Chipot C, Tajkhorshid E. 2020. Scalable molecular dynamics on CPU and GPU architectures with NAMD. J Chem Phys 153:044130.

46. Zhu X, Lopes PEM, Mackerell AD. 2012. Recent Developments and Applications of the CHARMM force fields. Wiley Interdiscip Rev Comput Mol Sci 2:167–185.

47. Schwarz DS, Blower MD. 2016. The endoplasmic reticulum: structure, function and response to cellular signaling. Cellular and Molecular Life Sciences 73:79–94.

48. English AR, Voeltz GK. 2013. Endoplasmic reticulum structure and interconnections with other organelles. Cold Spring Harb Perspect Biol 5:a013227.

49. Voeltz GK, Rolls MM, Rapoport TA. 2002. Structural organization of the endoplasmic reticulum. EMBO Rep 3:944–950.

50. Phillips MJ, Voeltz GK. 2016. Structure and function of ER membrane contact sites with other organelles. Nat Rev Mol Cell Biol 17:69–82.

51. Chen S, Novick P, Ferro-Novick S. 2013. ER structure and function. Curr Opin Cell Biol 25:428–433.

52. Evans AS, Lennemann NJ, Coyne CB. 2020. BPIFB3 Regulates Endoplasmic Reticulum Morphology To Facilitate Flavivirus Replication. J Virol 94.

53. Gillespie LK, Hoenen A, Morgan G, Mackenzie JM. 2010. The Endoplasmic Reticulum Provides the Membrane Platform for Biogenesis of the Flavivirus Replication Complex. J Virol 84:10438–10447.

54. Kaufusi PH, Kelley JF, Yanagihara R, Nerurkar VR. 2014. Induction of Endoplasmic Reticulum-Derived Replication-Competent Membrane Structures by West Nile Virus Non-Structural Protein 4B. PLoS One 9:e84040.

55. Ngo AM, Shurtleff MJ, Popova KD, Kulsuptrakul J, Weissman JS, Puschnik AS. 2019. The ER membrane protein complex is required to ensure correct topology and stable expression of flavivirus polyproteins. Elife 8.

56. Paul D. 2013. Architecture and biogenesis of plus-strand RNA virus replication factories. World J Virol 2:32.

57. Paul D, Bartenschlager R. 2015. Flaviviridae Replication Organelles: Oh, What a Tangled Web We Weave. Annu Rev Virol 2:289–310.

58. Rajah MM, Monel B, Schwartz O. 2020. The entanglement between flaviviruses and ER-shaping proteins. PLoS Pathog 16:e1008389.

59. Lin DL, Inoue T, Chen Y-J, Chang A, Tsai B, Tai AW. 2019. The ER Membrane Protein Complex Promotes Biogenesis of Dengue and Zika Virus Non-structural Multi-pass Transmembrane Proteins to Support Infection. Cell Rep 27:1666–1674.e4.

60. Chambers TJ, Grakoui A, Rice CM. 1991. Processing of the yellow fever virus nonstructural polyprotein: a catalytically active NS3 proteinase domain and NS2B are required for cleavages at dibasic sites. J Virol 65:6042–6050.

61. Wahaab A, Liu K, Hameed M, Anwar MN, Kang L, Li C, Ma X, Wajid A, Yang Y, Khan UH, Wei J, Li B, Shao D, Qiu Y, Ma Z. 2021. Identification of Cleavage Sites Proteolytically Processed by NS2B-NS3 Protease in Polyprotein of Japanese Encephalitis Virus. Pathogens 10:102.

62. Bozzacco L, Yi Z, Andreo U, Conklin CR, Li MMH, Rice CM, MacDonald MR. 2016. Chaperone-Assisted Protein Folding Is Critical for Yellow Fever Virus NS3/4A Cleavage and Replication. J Virol 90:3212–3228.

63. Heaton NS, Perera R, Berger KL, Khadka S, LaCount DJ, Kuhn RJ, Randall G. 2010. Dengue virus nonstructural protein 3 redistributes fatty acid synthase to sites of viral replication and increases cellular fatty acid synthesis. Proceedings of the National Academy of Sciences 107:17345–17350.

64. Murray CL, Jones CT, Rice CM. 2008. Architects of assembly: roles of Flaviviridae non-structural proteins in virion morphogenesis. Nat Rev Microbiol 6:699–708.

65. Li Q, Kang C. 2022. Structures and Dynamics of Dengue Virus Nonstructural Membrane Proteins. Membranes (Basel) 12:231.

66. Lescar J, Soh S, Lee LT, Vasudevan SG, Kang C, Lim SP. 2018. The Dengue Virus Replication Complex: From RNA Replication to Protein-Protein Interactions to Evasion of Innate Immunity, p. 115–129. *In*.

67. Perera R, Kuhn RJ. 2008. Structural proteomics of dengue virus. Curr Opin Microbiol 11:369–77.

68. Constant DA, Mateo R, Nagamine CM, Kirkegaard K. 2018. Targeting intramolecular proteinase NS2B/3 cleavages for trans -dominant inhibition of dengue virus. Proceedings of the National Academy of Sciences 115:10136–10141.

69. Luo D, Xu T, Hunke C, Grüber G, Vasudevan SG, Lescar J. 2008. Crystal Structure of the NS3 Protease-Helicase from Dengue Virus. J Virol 82:173–183.

70. Zhang L, Mohan PM, Padmanabhan R. 1992. Processing and localization of Dengue virus type 2 polyprotein precursor NS3-NS4A-NS4B-NS5. J Virol 66:7549–7554.

71. Preugschat F, Yao CW, Strauss JH. 1990. In vitro processing of dengue virus type 2 nonstructural proteins NS2A, NS2B, and NS3. J Virol 64:4364–4374.

72. Welsch S, Miller S, Romero-Brey I, Merz A, Bleck CKE, Walther P, Fuller SD, Antony C, Krijnse-Locker J, Bartenschlager R. 2009. Composition and Three-Dimensional Architecture of the Dengue Virus Replication and Assembly Sites. Cell Host Microbe 5:365–375.

73. Cortese M, Goellner S, Acosta EG, Neufeldt CJ, Oleksiuk O, Lampe M, Haselmann U, Funaya C, Schieber N, Ronchi P, Schorb M, Pruunsild P, Schwab Y, Chatel-Chaix L, Ruggieri A, Bartenschlager R. 2017. Ultrastructural Characterization of Zika Virus Replication Factories. Cell Rep 18:2113–2123.

74. Preugschat F, Lenches EM, Strauss JH. 1991. Flavivirus enzyme-substrate interactions studied with chimeric proteinases: identification of an intragenic locus important for substrate recognition. J Virol 65:4749–4758.

75. Erbel P, Schiering N, D’Arcy A, Renatus M, Kroemer M, Lim SP, Yin Z, Keller TH, Vasudevan SG, Hommel U. 2006. Structural basis for the activation of flaviviral NS3 proteases from dengue and West Nile virus. Nat Struct Mol Biol 13:372–373.

76. Chappell KJ, Nall TA, Stoermer MJ, Fang N-X, Tyndall JDA, Fairlie DP, Young PR. 2005. Site-directed Mutagenesis and Kinetic Studies of the West Nile Virus NS3 Protease Identify Key Enzyme-Substrate Interactions. Journal of Biological Chemistry 280:2896– 2903.

77. Li J, Lim SP, Beer D, Patel V, Wen D, Tumanut C, Tully DC, Williams JA, Jiricek J, Priestle JP, Harris JL, Vasudevan SG. 2005. Functional Profiling of Recombinant NS3 Proteases from All Four Serotypes of Dengue Virus Using Tetrapeptide and Octapeptide Substrate Libraries. Journal of Biological Chemistry 280:28766–28774.

78. Roosendaal J, Westaway EG, Khromykh A, Mackenzie JM. 2006. Regulated cleavages at the West Nile virus NS4A-2K-NS4B junctions play a major role in rearranging cytoplasmic membranes and Golgi trafficking of the NS4A protein. J Virol 80:4623–32.

79. Amberg SM, Rice CM. 1999. Mutagenesis of the NS2B-NS3-Mediated Cleavage Site in the Flavivirus Capsid Protein Demonstrates a Requirement for Coordinated Processing. J Virol 73:8083–8094.

80. Lin C, Amberg SM, Chambers TJ, Rice CM. 1993. Cleavage at a novel site in the NS4A region by the yellow fever virus NS2B-3 proteinase is a prerequisite for processing at the downstream 4A/4B signalase site. J Virol 67:2327–2335.

81. Franz KM, Neidermyer WJ, Tan Y-J, Whelan SPJ, Kagan JC. 2018. STING-dependent translation inhibition restricts RNA virus replication. Proceedings of the National Academy of Sciences 115:E2058–E2067.

82. Liu Y, Gordesky-Gold B, Leney-Greene M, Weinbren NL, Tudor M, Cherry S. 2018. Inflammation-Induced, STING-Dependent Autophagy Restricts Zika Virus Infection in the Drosophila Brain. Cell Host Microbe 24:57–68.e3.

83. McGuckin Wuertz K, Treuting PM, Hemann EA, Esser-Nobis K, Snyder AG, Graham JB, Daniels BP, Wilkins C, Snyder JM, Voss KM, Oberst A, Lund J, Gale M. 2019. STING is required for host defense against neuropathological West Nile virus infection. PLoS Pathog 15:e1007899.

84. Aguirre S, Maestre AM, Pagni S, Patel JR, Savage T, Gutman D, Maringer K, Bernal-Rubio D, Shabman RS, Simon V, Rodriguez-Madoz JR, Mulder LCF, Barber GN, Fernandez-Sesma A. 2012. DENV Inhibits Type I IFN Production in Infected Cells by Cleaving Human STING. PLoS Pathog 8:e1002934.

85. Su C-I, Kao Y-T, Chang C-C, Chang Y, Ho T-S, Sun HS, Lin Y-L, Lai MMC, Liu Y-H, Yu C-Y. 2020. DNA-induced 2′3′-cGAMP enhances haplotype-specific human STING cleavage by dengue protease. Proceedings of the National Academy of Sciences 117:15947– 15954.

86. Ding Q, Gaska JM, Douam F, Wei L, Kim D, Balev M, Heller B, Ploss A. 2018. Species-specific disruption of STING-dependent antiviral cellular defenses by the Zika virus NS2B3 protease. Proceedings of the National Academy of Sciences 115.

87. Luo X, Yuan Y, Ma X, Luo X, Chen J, Chen C, Yang X, Yang J, Zhu X, Li M, Liu Y, Zhang P, Liu C. 2024. Diacylglycerol O-acyltransferase 2, a Novel Target of Flavivirus NS2B3 Protease, Promotes Zika Virus Replication by Regulating Lipid Droplet Formation. Research (Wash D C) 7:0511.

88. Martins AS, Martins IC, Santos NC. 2018. Methods for Lipid Droplet Biophysical Characterization in Flaviviridae Infections. Front Microbiol 9.

89. Cloherty APM, Olmstead AD, Ribeiro CMS, Jean F. 2020. Hijacking of Lipid Droplets by Hepatitis C, Dengue and Zika Viruses—From Viral Protein Moonlighting to Extracellular Release. Int J Mol Sci 21:7901.

90. Qin Z-L, Yao Q-F, Ren H, Zhao P, Qi Z-T. 2022. Lipid Droplets and Their Participation in Zika Virus Infection. Int J Mol Sci 23:12584.

91. Gandikota C, Mohammed F, Gandhi L, Maisnam D, Mattam U, Rathore D, Chatterjee A, Mallick K, Billoria A, Prasad VS V., Sepuri NBV, Venkataramana M. 2020. Mitochondrial Import of Dengue Virus NS3 Protease and Cleavage of GrpEL1, a Cochaperone of Mitochondrial Hsp70. J Virol 94.

92. Yu C-Y, Liang J-J, Li J-K, Lee Y-L, Chang B-L, Su C-I, Huang W-J, Lai MMC, Lin Y-L. 2015. Dengue Virus Impairs Mitochondrial Fusion by Cleaving Mitofusins. PLoS Pathog 11:e1005350.

93. Junjhon J, Pennington JG, Edwards TJ, Perera R, Lanman J, Kuhn RJ. 2014. Ultrastructural characterization and three-dimensional architecture of replication sites in dengue virus-infected mosquito cells. J Virol 88:4687–97.

94. Płaszczyca A, Scaturro P, Neufeldt CJ, Cortese M, Cerikan B, Ferla S, Brancale A, Pichlmair A, Bartenschlager R. 2019. A novel interaction between dengue virus nonstructural protein 1 and the NS4A-2K-4B precursor is required for viral RNA replication but not for formation of the membranous replication organelle. PLoS Pathog 15:e1007736.

95. Cortese M, Mulder K, Chatel-Chaix L, Scaturro P, Cerikan B, Plaszczyca A, Haselmann U, Bartenschlager M, Neufeldt CJ, Bartenschlager R. 2021. Determinants in Nonstructural Protein 4A of Dengue Virus Required for RNA Replication and Replication Organelle Biogenesis. J Virol 95.

96. Kiemel D, Kroell A-SH, Denolly S, Haselmann U, Bonfanti J-F, Andres JI, Ghosh B, Geluykens P, Kaptein SJF, Wilken L, Scaturro P, Neyts J, Van Loock M, Goethals O, Bartenschlager R. 2024. Pan-serotype dengue virus inhibitor JNJ-A07 targets NS4A-2K-NS4B interaction with NS2B/NS3 and blocks replication organelle formation. Nat Commun 15:6080.

97. Huang L, Pike D, Sleat DE, Nanda V, Lobel P. 2014. Potential Pitfalls and Solutions for Use of Fluorescent Fusion Proteins to Study the Lysosome. PLoS One 9:e88893.

